# Simultaneous optimization of lignocellulosic sugar catabolism via systematic laboratory evolution in dynamic conditions

**DOI:** 10.64898/2026.02.02.702459

**Authors:** Sunghwa Woo, Hyun Gyu Lim, Brenna Norton-Baker, Torrey M. Lind, Nathan E. Gladden, Yan Chen, Thomas Eng, Christopher W. Johnson, Aindrila Mukhopadhyay, Christopher J. Petzold, Adam M. Guss, Gregg T. Beckham, Adam M. Feist

**Author notes:** These authors made equal contributions to this work (Sunghwa Woo, Hyun Gyu Lim).

## Abstract

Efficient co-utilization of hexose and pentose sugars from lignocellulose is essential for microbial production of bio-based chemicals, yet engineered non-native catabolic pathways can be suboptimal and evolutionarily unstable in complex resource environments. We used a *Pseudomonas putida* strain, previously engineered to catabolize xylose and arabinose to examine how resource abundance, temporal availability, and sub-culturing criteria shape evolutionary outcomes. Using an automated adaptive laboratory evolution (ALE) platform, we evolved the strain under static conditions with single selection pressures and dynamic regimes that imposed selection pressures on multiple sugars. These environments drove divergence between catabolic specialists and generalists. While selection regimes with weak or absent selection for xylose frequently resulted in loss of xylose catabolism, evolution under carbon-limited, mixed-sugar environments promoted stable retention and coordinated optimization of multiple catabolic pathways, increasing total sugar consumption in mixed-sugar conditions. Genomic, proteomic, and biochemical analyses showed that evolutionary stability was determined by pathway-specific fitness costs, leading to either pathway loss or cost-reducing refinement, depending on selection strength. An isolated generalist clone also exhibited improved indigoidine production from mixed sugars when compared to the parental strain. Together, these findings link resource dynamics to fitness landscapes that determine catabolic specialization, generalization, evolutionary trade-offs, and applicability to bioconversion.

## Introduction

A major challenge in lignocellulosic bioconversion is enabling microbes to efficiently assimilate mixed sugars under industrially relevant conditions. Lignocellulosic hydrolysates contain both hexoses and pentoses, and biocatalysts must co-utilize these substrates to maximize production yields and rates^1–4^. Recent advances in industrial biotechnology have enhanced the efficiency of microbial biocatalysts for converting biomass feedstock into value-added chemical products^5,6^. Nevertheless, the engineering of such microbes is complex, given the regulatory and metabolic strategies that microbes have evolved to handle fluctuating substrate availability and ecological competition in natural environments^7–9^. As a result, microbial species often exhibit distinct substrate preferences, and hierarchical substrate utilization is governed by mechanisms such as carbon catabolite repression, which promotes rapid growth and competitive dominance by prioritizing preferred substrates^10–12^. While advantageous in natural ecosystems, such regulatory mechanisms hinder the simultaneous assimilation of mixed substrates, which is a critical limitation for microbial biocatalysts designed to process biomass-derived feedstocks^13–15^.

*Pseudomonas putida* has emerged as an attractive microbial host for bioproduction, owing to its stress tolerance and ability to catabolize aromatic compounds^16–18^. Engineering efforts have expanded its substrate range through the introduction of multiple variants of enabling catabolic pathways^19–26^. Building on these advances, the JE3692 strain enabled simultaneous utilization of multiple substrates, including glucose, xylose, arabinose, acetate, and *p*-coumarate, via chromosomal integrated pentose catabolic pathways, adaptive laboratory evolution (ALE), and alleviation of carbon catabolite repression^23^. Despite this progress, fully optimizing the simultaneous utilization of multiple sugars, especially non-native substrates, remains difficult due to complex regulatory and metabolic interplay.

ALE is a powerful approach to strain optimization that can help overcome strain inefficiencies via diversity generation and selection^27^. Most ALE studies to date have been conducted under static conditions with a single selection pressure^28–33^. Such regimes may not impose sufficient or appropriate selection pressure for performance in mixed-sugar contexts, as catabolic pathways not under direct selection often remain unoptimized. Moreover, heterologous or engineered pathways that reduce fitness can be lost, unless their expression is directly coupled to survival. While some studies have attempted co-optimization under multiple selection pressures within a single ALE campaign, outcomes have typically included either limited fitness improvements compared to single-substrate ALE or the loss of substrate-specific beneficial mutations^34,35^. This underscores the need for alternative ALE strategies that incorporate environmental complexity and dynamics to drive balanced optimization of multiple substrates.

Here, we used an automated ALE platform to evolve the previously engineered *P. putida* strain, JE3692, carrying heterologous xylose and arabinose catabolic pathways^23^ under both single-sugar and mixed-sugar regimes. This comparative approach allowed us to probe how environmental composition and temporal dynamics shape evolutionary trajectories in an application-driven context. Multi-omics analyses revealed distinct adaptive strategies, including regulatory rewiring and selective pathway retention. Furthermore, a strain with improved mixed-sugar utilization exhibited enhanced production of the non-ribosomal peptide indigoidine, underscoring the biotechnological potential of evolution-guided strategies. Our findings demonstrate that dynamic ALE can be used to dissect carbon catabolism by revealing how differential substrate utilization rates, pathway-specific fitness costs, and time-varying selection pressures shape evolutionary outcomes, providing design guidelines for engineering microbial platforms for mixed and heterogeneous feedstocks as well as similar complex evolutionary engineering goals.

## Results

### Systematic dynamic evolution for optimized utilization of the major lignocellulosic sugars

The engineered *Pseudomonas putida* strain JE3692 was used as the starting strain for all ALE campaigns^23^. This strain natively utilizes glucose and was previously modified to catabolize xylose and arabinose, two additional major sugars commonly derived from hemicellulose (**Figure 1A**). This strain harbors two heterologous catabolic pathways: for xylose utilization, the xylose isomerase pathway from *Escherichia coli* includes *xylE*, *xylA*, *xylB*, *tkt*, and *tal*. For arabinose metabolism, the oxidative arabinose pathway includes *araE1* from *E. coli* and *araA2*, *araB2*, *araC2*, *araD2*, and *araE2* from *Burkholderia ambifaria* AMMD^36^. In JE3692 strain, two deletions were introduced: *gcd* (glucose dehydrogenase) was deleted to prevent the formation of gluconate and xylonate^21,37^, and *crc* (catabolite repression control protein) was deleted to avoid the native regulation for sugar catabolism^14,25,38^. This strain represented both a promising candidate to build upon for industrial bioprocessing and a relevant test case for exploring dynamic ALE-driven optimization given its combination of native and engineered metabolic capabilities.

**Figure 1.**
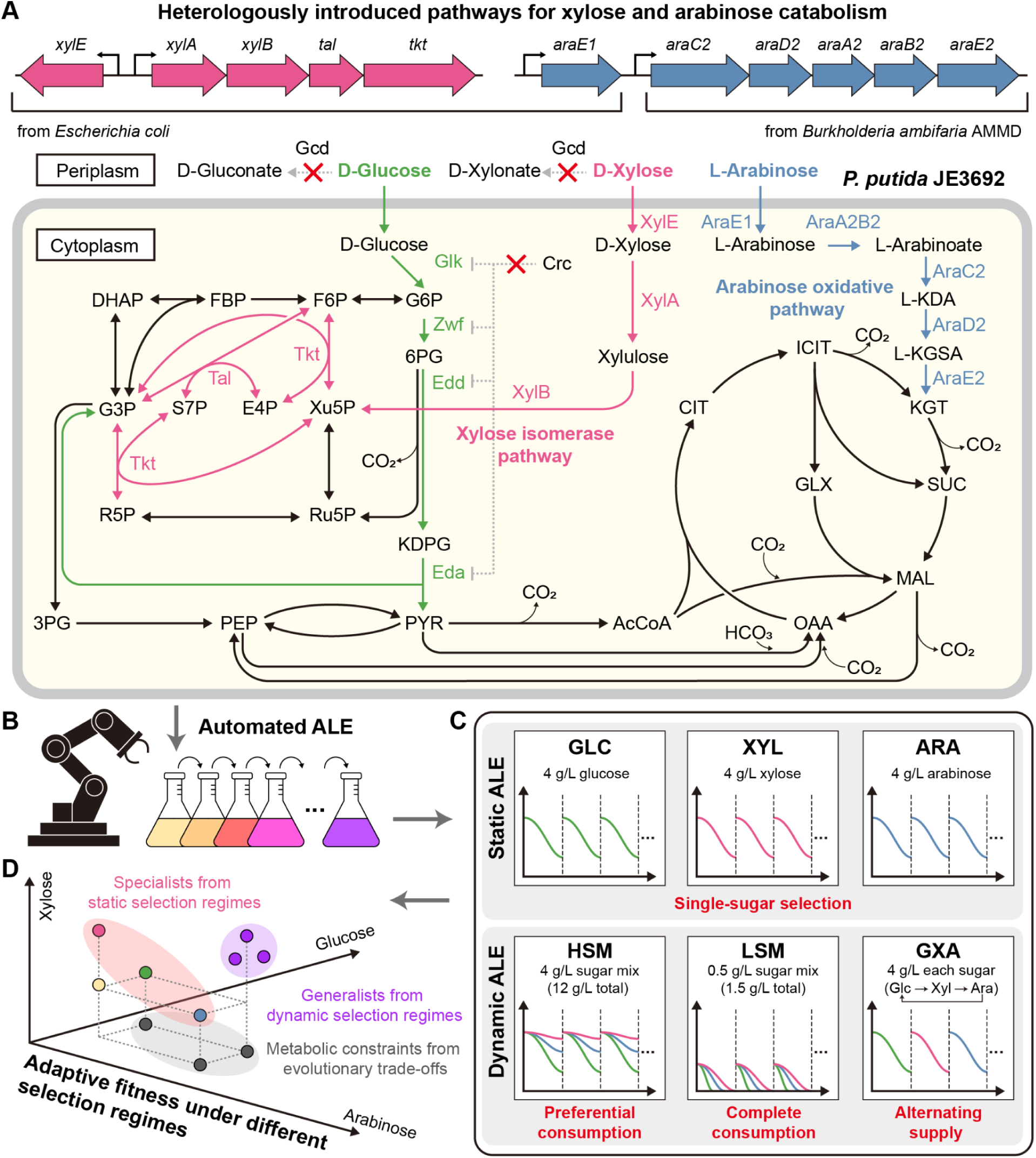
Systematic dynamic evolution for optimized utilization of multiple sugars. **(A)** Catabolic pathways for glucose, xylose, and arabinose in JE3692. Genes for each pathway are highlighted with a different color: green, endogenous Entner-Doudoroff (ED) pathway; pink, heterologous xylose isomerase pathway; blue, heterologous arabinose oxidative pathway. Gray dotted lines and red crosses indicate the deleted genes (*gcd* and *crc*). Abbreviations: G6P, D-glucose 6-phosphate; 6PG, D-gluconate 6-phosphate, KDPG, 2-dehydro-3-deoxy-D-gluconate 6-phosphate; F6P, D-fructose 6-phosphate; FBP, D-fructose 1,6-bisphosphate; DHAP, dihydroxyacetone phosphate; G3P, D-glyceraldehyde 3-phosphate; 3PG, 3-phospho-D-glycerate; PEP, phosphoenolpyruvate; PYR, pyruvate; AcCoA, acetyl-CoA, CIT, citrate; ICIT, isocitrate; KGT, 2-ketoglutarate; SUC, succinate; MAL, malate; OAA, oxaloacetate; GLX, glyoxylate; X5P, xylulose 5-phosphate; Ru5P, ribulose 5-phosphate; R5P, ribose 5-phosphate; E4P, D-erythrose 4-phosphate; S7P, D-sedoheptulose 7-phosphate; L-KDA, 2-keto-3-deoxy-L-arabinose; L-KGSA, 2-ketoglutarate semialdehyde. **(B)** Automated ALEbot robotic platform, which enabled consistent, systematic, and parallel execution of multiple independently controlled evolution lineages, thereby maximizing consistency. **(C)** ALE strategies used in this study. Static ALE groups: GLC (glucose 4 g/L), XYL (xylose 4 g/L), ARA (arabinose 4 g/L). Dynamic ALE groups: HSM (glucose 4 g/L, xylose 4 g/L, and arabinose 4 g/L; selective sugar consumption allowed), LSM (glucose 0.5 g/L, xylose 0.5 g/L, and arabinose 0.5 g/L; complete sugar consumption mandated), GXA (glucose 4 g/L, xylose 4 g/L, or arabinose 4 g/L; each sugar alternatively supplied in series). Each ALE was performed with four replicates (*n* = 4). The differently colored curves in the simplified graph represent the expected sugar consumption profiles for each passage, separated by dotted lines: green for glucose, pink for xylose, and blue for arabinose. **(D)** A conceptual diagram of the fitness landscape highlighting potential evolutionary outcomes. Selected phenotypes may range from catabolic specialists, optimized for catabolism of a specific substrate but often incurring trade-offs on others, to catabolic generalists, which utilize a range of targeted metabolites without significant trade-offs on any one substrate.

The baseline fitness of JE3692 was evaluated in single-sugar and mixed-sugar conditions. The strain exhibited a specific growth rate of 0.42 ± 0.04 h^-1^ in glucose minimal medium, while growth rates in xylose and arabinose minimal media were lower, at 0.20 ± 0.01 and 0.14 ± 0.02 h^-1^, respectively. In a mixed-sugar minimal medium (4 g/L each), the strain grew at 0.46 ± 0.04 h^-1^, 23% slower than the previously reported rate (0.60 h^-1^)^23^. These lower growth rates on xylose and arabinose relative to glucose highlight differences in catabolic efficiency among the pathways and motivate further optimization of the heterologous sugar utilization systems^25^.

To investigate variation in evolutionary adaptation across different sugar conditions and to enable robust, multi-sugar conversion^34^, we performed multiple ALE experiments under both static and dynamic conditions. All ALE experiments were conducted using an automated robotic ALEbot platform, which enabled consistent, systematic, and parallel execution of multiple, independently-controlled evolution lines, thereby minimizing human error^39^ (**Figure 1B**). The ALE trajectories were categorized into static and dynamic selection conditions (**Figure 1C**). In static ALE, modified M9 minimal medium was supplemented with 4 g/L of a single sugar, namely glucose (GLC), xylose (XYL), or arabinose (ARA). Dynamic ALE consisted of three regimes: HSM (4 g/L each glucose, xylose, and arabinose) with OD-based transfer; LSM (0.5 g/L each) with sugar depletion-based transfer; and GXA, namely sequential exposure to glucose, xylose, and arabinose (4 g/L each). While GXA introduces dynamic selection through explicit sequential sugar exposure, the mixed-sugar conditions (HSM and LSM) are also dynamic in that differential utilization rates of glucose, xylose, and arabinose lead to temporal shifts in substrate availability and selection pressure within each growth cycle. Each ALE condition was performed in quadruplicate, yielding 24 independent lineages.

ALE experiments were executed until growth improvements plateaued in at least three of the four replicates per condition. Across the approximately 3-month experiments, cultures underwent 63–109 passages, corresponding to 394–709 generations and 32.4–61.5 × 10^11^ cumulative cell divisions (CCDs) (**Supplementary Table 1**)^40^. Growth rate improvements were typically observed early and tended to stabilize after 34–55 passages (**Supplementary Figure 1**). Clonal isolates were obtained from all end-point populations for phenotypic and genotypic characterization, as they were expected to exhibit distinct captive fitness depending on their respective selection regimes (**Figure 1D**).

### Static ALE selected for substrate-specific catabolic adaptation and trade-offs in unselected functions

The specific growth rates of evolved isolates from the static ALEs were compared to assess their fitness improvements on each sugar. The GLC isolates showed an insignificant increase in growth rate in glucose minimal medium (*p* > 0.05) (**Figure 2A**). The highest growth rate was 0.56 h^-1^, about 66% of that reported for evolved wild-type *P. putida* populations (0.85 h^-1^)^41^, likely reflecting disruption of the gluconate branch of glucose catabolism (*gcd* gene deletion). The XYL and ARA isolates exhibited improved growth when cultured in their respective selection conditions. Specifically, the growth rates improved by up to 1.5-fold for xylose (0.29 h^-1^) and 4.1-fold for arabinose (0.60 h^-1^) (**Figure 2B-C**), suggesting that adaptive mutations in the corresponding sugar catabolic pathways were acquired during evolution.

**Figure 2.**
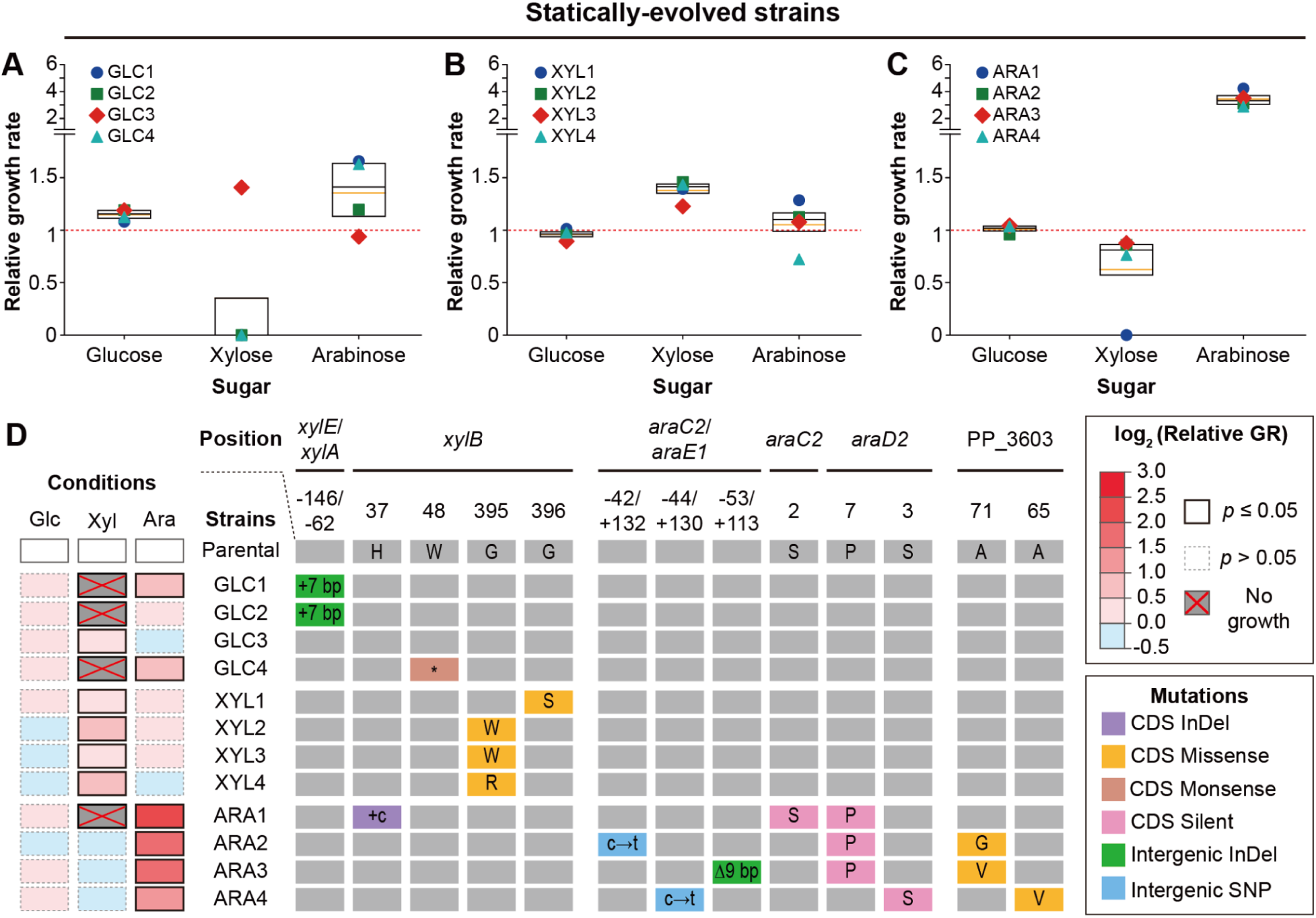
Sugar-specific adaptations and trade-offs in statically evolved strains. Box plots show the relative growth rates of statically evolved strains isolated from **(A)** GLC, **(B)** XYL, and **(C)** ARA conditions under single-sugar conditions (glucose, xylose, and arabinose), compared to the starting JE3692 strain. Each dot represents the mean fold change in specific growth rate from duplicates (*n* = 2). Each box indicates the interquartile range, and each black and orange solid line in the box indicates the median and mean value, respectively. Red dotted lines indicate a fold change of 1. The *x*-axis represents sugar conditions under which the isolates were grown, and the *y*-axis represents the specific growth rate (h^-1^). **(D)** Detailed mutational analysis related to xylose and arabinose catabolism in statically evolved strains. The right side shows mutations in the heterologous xylose and arabinose catabolic pathways. Rows and columns represent isolates and mutational positions, respectively. Uppercase and lowercase letters in boxes indicate amino acids and nucleobases, respectively. Different colors represent different mutation types detailed in the legend. The heatmap on the left shows the log fold change in the specific growth rate of each evolved isolate compared to the parental strain under various sugar conditions, generated from the same dataset as Figure 2A-C. The rows and columns represent isolate and sugar conditions under which the isolates were grown, respectively. Boxes with the color scale represent a gradient according to the log fold change in the specific growth rate (relative GR), showing a range of minimum (light blue) to maximum (red), and gray boxes marked with cross indicate no growth.

Beyond condition-specific improvements, notable unexpected impacts on alternative sugar catabolism were also observed. Notably, three GLC (1, 2, 4) and one ARA (1) isolates failed to grow on xylose, suggesting an inactivation of the heterologous xylose pathway (*vide infra*, **Figure 2A, C**). Meanwhile, several GLC isolates exhibited enhanced growth on alternative sugars, particularly arabinose. Namely, the GLC1 and GLC4 strains showed elevated growth in arabinose medium, despite arabinose not being included in their original selection regime (**Figure 2A**). All statically evolved isolates showed no significant improvement in growth rate in HSM medium (4 g/L each of glucose, xylose, and arabinose), indicating that sugar-specific adaptations may not translate into improved fitness in mixed environments and motivating alternative ALE approaches (**Supplementary Figure 2**).

To identify genetic changes associated with the observed physiology, whole-genome sequencing was performed on the statically evolved isolate, revealing key mutations that enhanced fitness on xylose and arabinose, and no mutations were identified in the glucose catabolic pathway of the GLC1-4 strains, consistent with no significant improvement in growth rate in glucose medium. Recurrent mutations were observed within each lineage: the XYL strains consistently carried mutations in only *xylB*, whereas the Ara strains shared mutations in *araD2*, *araC2*, *araE1*, or their intergenic region (**Figure 2D**). Additionally, a native gene, PP_3603 encoding a transcriptional regulator of the glucuronate and glucarate catabolic pathways, was also mutated across ARA isolates, suggesting its involvement in arabinose catabolism.

We also identified endogenous genes that were repeatedly mutated across multiple statically evolved lineages, indicating convergent adaptation. Of the 28 mutations identified in regions repeatedly mutated across independent lineages, 12 mapped to functional genes, including PP_1650 (GacS, a two-component histidine kinase), PP_4171 (hypothetical protein), PP_4173 (two component sensor histidine kinase/response regulator), and PP_4373 (FleQ, a transcriptional regulator) (**Supplementary Figure 3**). We interpret these changes as generally beneficial adaptations to laboratory environments, based on their convergence across multiple ALE groups and observed occurrence in previous ALE studies with *P. putida* KT2440^41–43^.

The *xylB* gene emerged as a common target wherein mutations either improved or impaired cellular fitness on xylose (**Supplementary Figure 4**). Mutations in *xylB* of the XYL1-4 strains occurred at amino acids 395 and 396 (**Figure 2D**), which are highly conserved residues in XylB homologues (**Supplementary Figure 5)**. Disruptive *xylB* mutations included a frameshift (GLC4) and a premature stop codon (ARA1). The other isolates incapable of xylose catabolism exhibited an identical insertion mutation in the intergenic region between *xylE* and *xylA*, potentially reducing expression of the downstream xylose transporter and catabolic enzymes. Interestingly, these two divergent mutational strategies in the xylose pathway revealed the emergence of catabolic specialists and the trade-offs associated with a functional loss not under selection during the evolution experiments.

Interestingly, compared to the xylose pathway, no disruptive mutations were found in the arabinose pathway (**Figure 2D**). This distinction is important because both pathways are heterologous, yet only one appears to be more favored. All 8 observed mutations in this pathway were found exclusively in ARA isolates: 3 occurred in the intergenic region upstream of the *araCDAB2* operon, and 5 were in the N-terminal region of the *araC2* or *araD2* genes, all which were likely involved in regulating expression levels^44^. Strains with these mutations exhibited significantly increased growth rates in arabinose minimal media. Additionally, 3 mutations in the PP_3603 gene likely enhanced arabinose catabolism by modulating the expression of PP_3602, which shares the same functional annotation as the *araE2* gene (2-ketoglutarate semialdehyde dehydrogenase)^45,46^. Collectively, the phenotyping and genomic analysis of the statically evolved strains showed trends of condition-specific optimization and generation of catabolic specialists with no significant improvement of growth on sugar mixtures (*p* > 0.05).

### Dynamic ALE revealed the selection of catabolic generalists dependent on experimental design

The dynamic ALE also enabled generation of both metabolic specialists and catabolic generalists, shaped by their specific selection regimes. As observed for the GLC and ARA statically evolved isolates, the HSM isolates, which did not require complete sugar consumption, selectively lost the ability to grow on xylose as the sole carbon source (**Figure 3A**), indicating that the xylose pathway is detrimental under these conditions. Two HSM isolates showed higher growth rates on arabinose alone whereas their growth rates on glucose alone remained similar to the unevolved parent. Conversely, the LSM and GXA isolates displayed enhanced growth rates on xylose, indicating xylose catabolism was beneficial in these conditions (**Figure 3B-C**). Evolution under the LSM condition also greatly enhanced the growth rate on arabinose, up to 3.0-fold, comparable to the growth rates of the ARA isolates. In contrast, the growth rates on arabinose alone of the GXA isolates were not significantly increased (**Figure 3C**).

**Figure 3.**
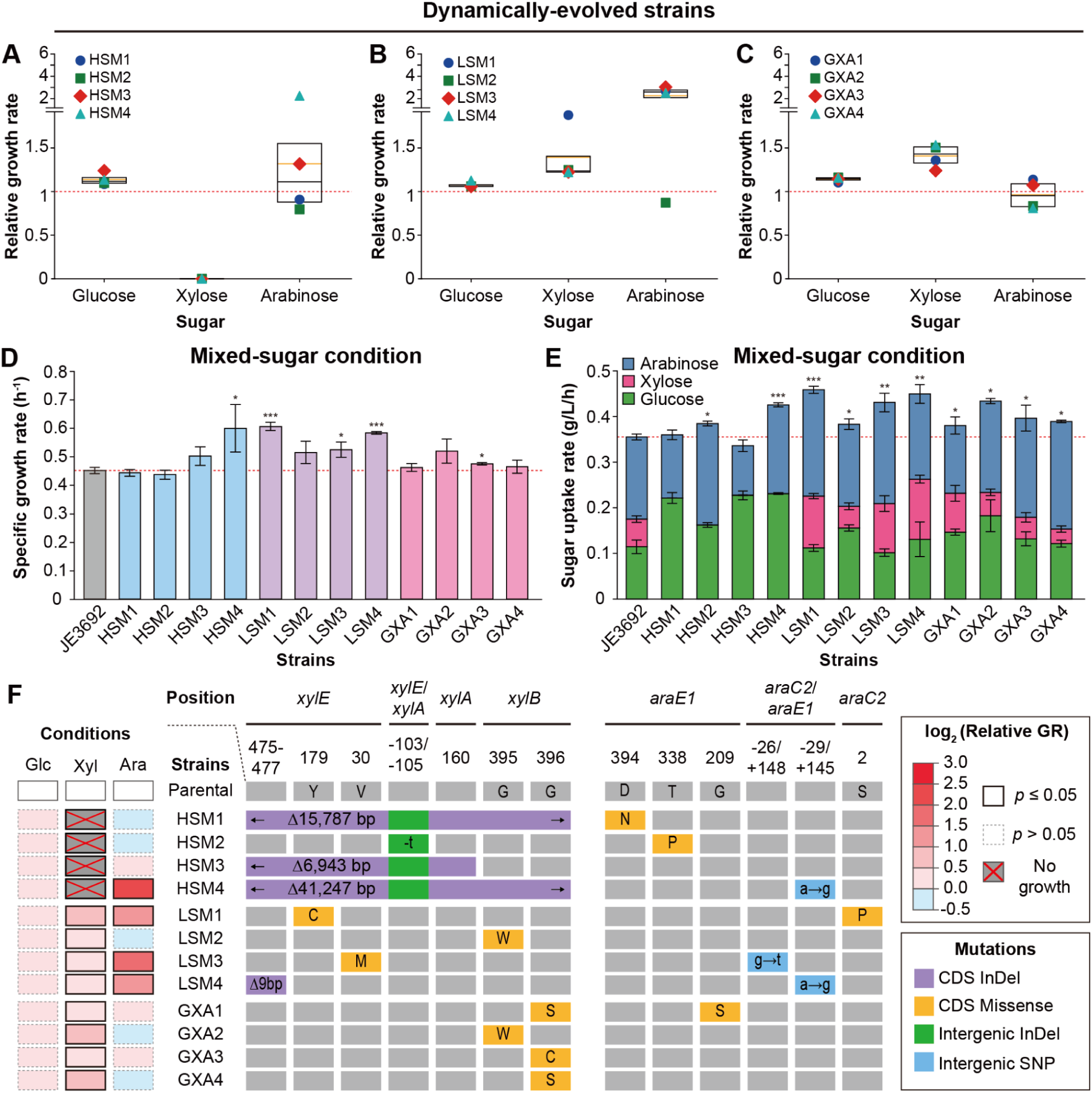
Dynamic selection resulted in the generation of catabolic generalists or strains with trade-offs dependent on media conditions. Box plots show the relative growth rates of dynamically evolved strains isolated from **(A)** HSM, **(B)** LSM, and **(C)** GXA conditions under single-sugar conditions (glucose, xylose, and arabinose), compared to the parental strain, JE3692. Each dot represents the mean fold change in specific growth rate from duplicates (*n* = 2). **(D)** Specific growth rates of dynamically evolved strains under the HSM mixed-sugar conditions. The *x*- and *y*-axes indicate strains and specific growth rate (h^-1^), respectively. The gray, light blue, lavender, and pink boxes represent JE3692, HSM, LSM, and GXA isolates, respectively. **(E)** Sugar uptake rates of dynamically evolved strains under the HSM mixed-sugar conditions. The *x*- and *y*-axes indicate strains and sugar uptake rate (g/L/h), respectively. The green, pink, and blue boxes represent glucose, xylose, and arabinose, respectively. Cultures were conducted in biological triplicates (*n* = 3). Red dotted lines indicate the specific growth rate and total sugar uptake rate of the JE3692 strain. Error bars indicate the standard deviations. *, **, and *** indicate *p* < 0.05, *p* < 0.01, and *p* < 0.001, respectively. Bars without asterisks represent values that are not statistically significantly different from those of the JE3692 strain. **(F)** Detailed mutational analysis related to xylose and arabinose catabolism in dynamically evolved strains. All other details are the same as those described for Figure 2D.

Growth and sugar consumption of the dynamically evolved strains were also evaluated under the mixed-sugar conditions containing 4 g/L each of glucose, xylose, and arabinose, identical to the HSM selection regime. Consistent with faster growth on individual sugars, the LSM isolates displayed generally higher growth rates than the other isolates. Among these, the LSM1 strain exhibited the highest growth rate (0.65 h^-1^), 1.4 times that of the JE3692 strain (**Figure 3D**). Moreover, during the exponential growth phase, the LSM isolates generally exhibited higher total sugar consumption rates than the other isolates (**Figure 3E**). Among them, the LSM1 strain showed the highest total sugar consumption rate of 0.46 g/L/h, which was 1.3-fold higher than that of the JE3692 strain. Its glucose consumption rate was comparable to that of JE3692, whereas the consumption rates for xylose and arabinose were 1.9- and 1.3-fold higher, respectively. Meanwhile, consistent with their inability to grow on xylose, the HSM isolates failed to consume xylose under mixed-sugar conditions. These results suggest that the requirement of complete consumption of all three sugars under the LSM condition was effective in simultaneously optimizing catabolism of lignocellulosic sugar mixes.

Genome resequencing of the dynamically evolved strains identified the mutations that occurred during evolution (**Figure 3F**). Firstly, a number of isolates carried mutations related to generally beneficial adaptation to laboratory environments, including mutations in PP_1650, PP_4171, PP_4173, and PP_4373, similar to those observed in statically evolved strains (**Supplementary Figure 3**). All the HSM isolates, which did not consume xylose, acquired large deletions in the *xylE*-*xylAB* region (HSM1, HSM3, HSM4) or an insertion upstream of *xylA* (HSM2), presumably leading to the observed elimination of xylose catabolism. In contrast, the LSM and GXA isolates, which retained the ability to utilize all three sugars, acquired non-disruptive mutations, defined here as sequence changes that did not result in loss-of-function, including substitutions or an in-frame deletion. In the arabinose pathway, mutations were identified in the protein sequences of *araE1* and *araC2*, as well as in the intergenic region upstream of *araC2*. However, mutations in *araE1* appeared to have no significant impact on fitness in arabinose medium (**Figure 3F**). The exclusive occurrence of disruptive mutations in the xylose pathway, but not in the arabinose pathway, suggested distinct fitness constraints and motivated the hypothesis that the xylose isomerase pathway imposes a higher fitness cost (*vide infra*) (**Supplementary Figure 3)**.

### Evolutionary stability of heterologous sugar catabolism under divergent selection regimes

To assess the causal effects of key non-disruptive ALE-derived mutations, we reverse-engineered several genetic changes back into the parent JE3692 strain and assessed fitness changes under xylose and arabinose conditions. In the xylose pathway, we selected *xylB* G395W, the most convergent mutation in the xylose pathway, and *xylE* Y179C, identified in the LSM1 strain. In the arabinose pathway, we selected *araC2* S2P, identified in the LSM1 strain, *araD2* P7P, the most convergent mutation in the arabinose pathway, and PP_3603 A71V, one of the mutations observed in the ARA isolate. Except for PP_3603 A71V, these mutations independently increased growth rates under the corresponding conditions (**Figure 4A**). Although introducing PP_3603 A71V alone reduced the growth rate on arabinose, it eliminated the initial lag phase, which may explain its persistence during the ALE experiments (**Supplementary Figure 6**). Moreover, evolved strains carrying mutations in PP_3603 (ARA 2–4) also harbored mutations in *araC2* or *araD2* (**Figure 2D**), suggesting that these mutations may modulate the effect of the PP_3603 A71V mutation on growth, although their synergistic effects are not clear.

**Figure 4.**
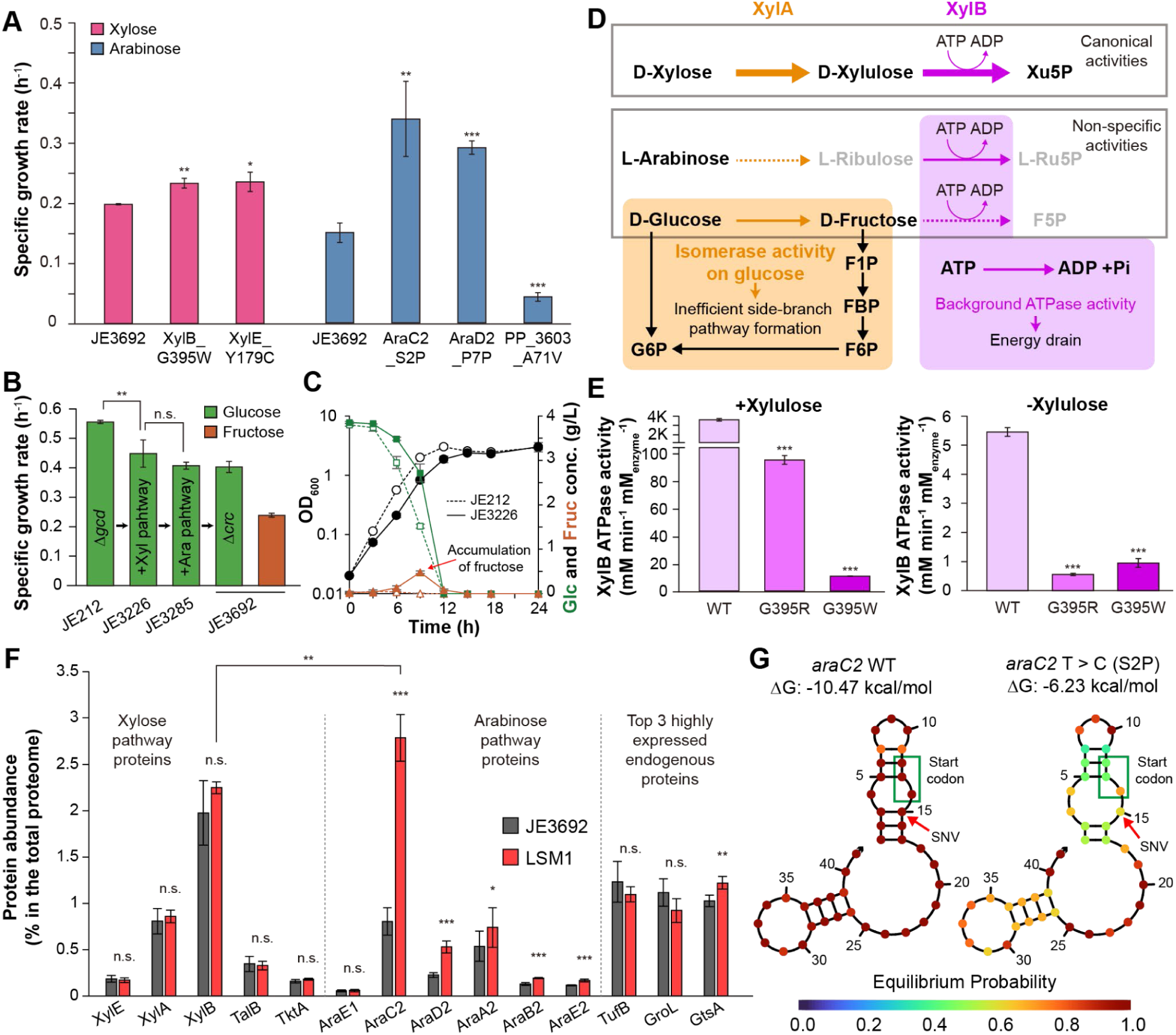
Evolution of heterologous sugar catabolism under divergent selective regimes. **(A)** Specific growth rates of JE3692 and reverse engineered strains cultured in xylose (pink) and arabinose (blue) minimal media. The *x*- and *y*-axes represent strains and specific growth rate (h^-1^), respectively. **(B)** Specific growth rates of the JE3692 and its intermediate strains cultured in glucose (green) and fructose (brown) minimal media. The *x*- and *y*-axes represent strains and specific growth rate (h^-1^), respectively. **(C)** Detailed culture profiles of JE212 (dotted lines and open symbols) and JE3226 (solid lines and closed symbols) in glucose minimal media. Data are from the same dataset as in panel B. The left and right *y*-axes indicate cell biomass (OD_­_) and concentration of glucose and fructose (g/L), respectively; the *x*-axis indicates time in hours (h). Symbols: black circles, OD_­_; green squares, glucose; brown triangles, fructose. **(D)** Overview of the proposed evolutionary scenario underlying the negative selection of xylose pathway enzymes. Orange and purple boxes and arrows indicate enzymatic activities of XylA and XylB, respectively. **(E)** *In vitro* comparison of ATPase activity between wild-type and mutant XylBs in the presence and absence of D-xylulose. Light, regular, and dark purple bars indicate wild type, G395R, and G395W, respectively. The *y*-axis represents ATPase activities (mM/min per mM enzyme), measured via NADH oxidation coupled to pyruvate kinase and lactate dehydrogenase reactions. **(F)** Comparison of the protein abundance levels of heterologous proteins for xylose and arabinose catabolism in JE3692 and LSM1. The *x*- and *y*-axes represent protein and the fraction in the total proteome, respectively. The gray and red boxes represent JE3692 and LSM1, respectively. **(G)** Comparison of secondary structure of 5’- end of mRNA of the wild-type and mutant *araC2*, predicted by NUPACK^55^. Modified base and start codon are highlighted by red circle and green box, respectively. Cultures and enzyme assay were conducted using biological triplicates (*n* = 3), and proteomic analysis was conducted using biological quadruplicates (*n* = 4). Error bars indicate the standard deviations. *, **, ***, and n.s. indicate *p* < 0.05, *p* < 0.01, *p* < 0.001, and *p* > 0.05, respectively. Asterisks without brackets indicate statistical significance compared to the parent.

In a separate assay, to understand the reason for the selective disruption of the xylose pathway during ALE, we examined whether the expression of the xylose and arabinose pathways imposed fitness costs under non-selective regimes using phenotypic assays. To this end, we cultivated the JE3692 and its intermediate strains in glucose minimal medium (**Figure 4B**). The strains utilized were JE212, which lacks both the xylose and arabinose pathways; JE3226, which contains only the xylose pathway; and JE3285, which contains both pathways. All three intermediate strains possessed an intact *crc* gene, which was deleted in the JE3692 strain. The JE3226 strain exhibited a 19% lower growth rate (0.45 ± 0.01 h^-1^, *p* < 0.01) compared to the JE212 strain (0.56 ± 0.05 h^-1^). Although the JE3285 strain showed a slightly lower growth rate (0.41 ± 0.01 h^-1^) than the JE3226 strain (a 9% decrease), the difference was not statistically significant (*p* > 0.05). We also observed a transient accumulation of up to 0.5 g/L fructose in the JE3226 culture grown in glucose minimal medium, whereas only negligible levels were detected in the JE212 culture (**Figure 4C**). Taken together, these results demonstrated that the xylose pathway reduced growth fitness under non-selective conditions, consistent with its selective loss during ALE in the absence of direct selection pressures.

To determine whether non-specific enzymatic activities of the xylose catabolic enzymes contributed to fitness costs and pathway instability during ALE^19,47–49^, we examined the biochemical activities of core enzymes, XylA and XylB, using *in vitro* enzyme assays (**Figure 4D**). In the XYL, LSM, and GXA groups, at least one isolate from each carried a substitution at residue 395 of XylB, and thus we compared the activities of the mutant XylB variants (G395R and G395W) (**Figure 2D** and **3F**). In *in vitro* enzyme assays, wild-type XylB exhibited high activity on D-xylulose and even retained background ATPase activity in the absence of a substrate (**Figure 4E**), consistent with previous studies^19,48^. Interestingly, the G395R and G395W variants exhibited markedly reduced xylulose activity, by 37-fold and 330-fold, respectively, despite having evolved under xylose-selective conditions. They also decreased background ATPase activity by 9.9- and 5.7-fold, respectively. Analysis of the AlphaFold-predicted structure suggested that residues 395 and 396 are positioned adjacent to the ATP-binding region (**Supplementary Figure 7**), suggesting that these substitutions may alter ATP binding affinity^50^. Although the background ATPase activity of wild-type XylB was only about 0.15% of that observed with D-xylulose, its high expression level may increase ATP turnover in the absence of xylose, potentially contributing to the impaired growth (**Figure 4B**). This outcome suggested that maximizing enzyme activity was not the primary determinant of fitness under selection. Accordingly, under strong selection pressure for xylose utilization (XYL, LSM, and GXA), XylB activity appeared to be fine-tuned while ATPase activity was reduced. In contrast, under weaker xylose selection pressure (GLC, ARA, and HSM), evolution tended to disrupt XylB or the entire xylose pathway.

The non-specific isomerase activity of XylA toward D-glucose may also influence fitness in this context. Proton NMR analysis qualitatively confirmed that XylA exhibited isomerase activity toward all three target sugars; high for D-xylose, moderate for D-glucose, and minimal for L-arabinose (**Supplementary Figure 8**)^47,49^. This activity toward glucose explains the transient accumulation of fructose in the JE3226 culture (**Figure 4C**). The formation of an inefficient side branch via unnecessary D-fructose production, accompanied by flux imbalance and diauxic sugar utilization^51,52^, may have imposed selective pressures to eliminate XylA during ALE experiments^53,54^. Further support comes from the observation that the JE3692 strain showed a 1.9-fold lower growth rate when grown on D-fructose than on D-glucose, likely due to the absence of phosphofructokinase (**Figure 4B**). This may explain why, in ALE regimes with strong selection pressure on glucose catabolism (GLC and HSM), not only XylB but the entire xylose pathway was disrupted by large deletions or mutations in intergenic regions.

In another drill-down assay on ALE-derived findings, proteomic analysis suggested that the high protein abundance of XylA and XylB may exacerbate the fitness burden imposed by the xylose isomerase pathway. XylB accounted for 2% and XylA for 0.8% of the total protein by mass (**Figure 4F**). Notably, the protein abundance of XylB was comparable to the total abundance of arabinose catabolic enzymes, and even exceeded that of the three most highly expressed endogenous proteins: TufB (elongation factor Tu-B), GroL (chaperonin), and GtsA (mannose/glucose ABC transporter substrate-binding protein) (**Supplementary Data 2**). These quantitative proteomic patterns indicate that high protein abundance of XylA and XylB amplified their non-specific activities and, in turn, the likely burdensome effects.

Finally, to investigate the arabinose pathway, we focused on the AraC2 S2P mutation and its associated proteomic changes in the LSM1 strain. Notably, comparative proteomic analysis revealed that AraC2 abundance in the LSM1 strain was 3.5-fold higher than in the JE3692 strain, due to the S2P mutation (**Figure 4F**). Furthermore, downstream arabinose catabolic enzymes, including AraD2, AraA2, AraB2, and AraE2, also showed increased protein abundance. These increases were likely attributable to a mutation near the 5′ end of *araC2* that destabilized the mRNA secondary structure around the ribosome-binding site, thereby improving ribosome accessibility and translation efficiency (**Figure 4G**)^55^. Indeed, based on UTR Designer predictions^44^, the mutant *araC2* was expected to show a 3.4-fold increase in expression.

Collectively, these findings demonstrated that evolutionary stability of heterologous pathways is determined by the balance between selection pressure and pathway-specific fitness costs. The arabinose pathway, which imposed minimal fitness burden, was stably retained across conditions. In contrast, the xylose pathway incurred substantial fitness costs and was therefore maintained only under regimes with strong selection for xylose utilization, where evolution favored mutations that reduced these costs rather than maximizing catalytic activity.

### An evolved isolate with a generalist sugar catabolism enhances indigoidine production

The potential of the evolutionarily optimized LSM1 strain evolved under dynamic conditions as a production host for conversion of lignocellulosic sugars was evaluated using indigoidine production, a natural blue pigment with various industrial applications, as a relevant bioproduct. To achieve this, we introduced the biosynthetic pathway for indigoidine into both the parental strain and the LSM1 strain (**Figure 5A**)^56–59^. This heterologous pathway included the *bpsA* gene encoding blue pigment synthetase A from *Streptomyces lavendulae* and the *sfp* gene encoding 4′-phosphopantetheinyl transferase from *Bacillus subtilis*^60,61^. Using the plasmid system, these two genes were expressed under the *tac* promoter without *lacI*, which has previously been used for strong and constitutive expression in *P. putida* KT2440^62,63^. This plasmid was transformed into JE3692 and LSM1, and the resulting strains, JE3692_IND and LSM1_IND, were cultivated in isotonic sugar media containing 4 g/L each of glucose, xylose, and arabinose, as well as in hydrolysate mimicking sugar media simulating lignocellulosic sugar composition, which contained 24 g/L of glucose, 12 g/L of xylose, and 4 g/L of arabinose^64^.

**Figure 5.**
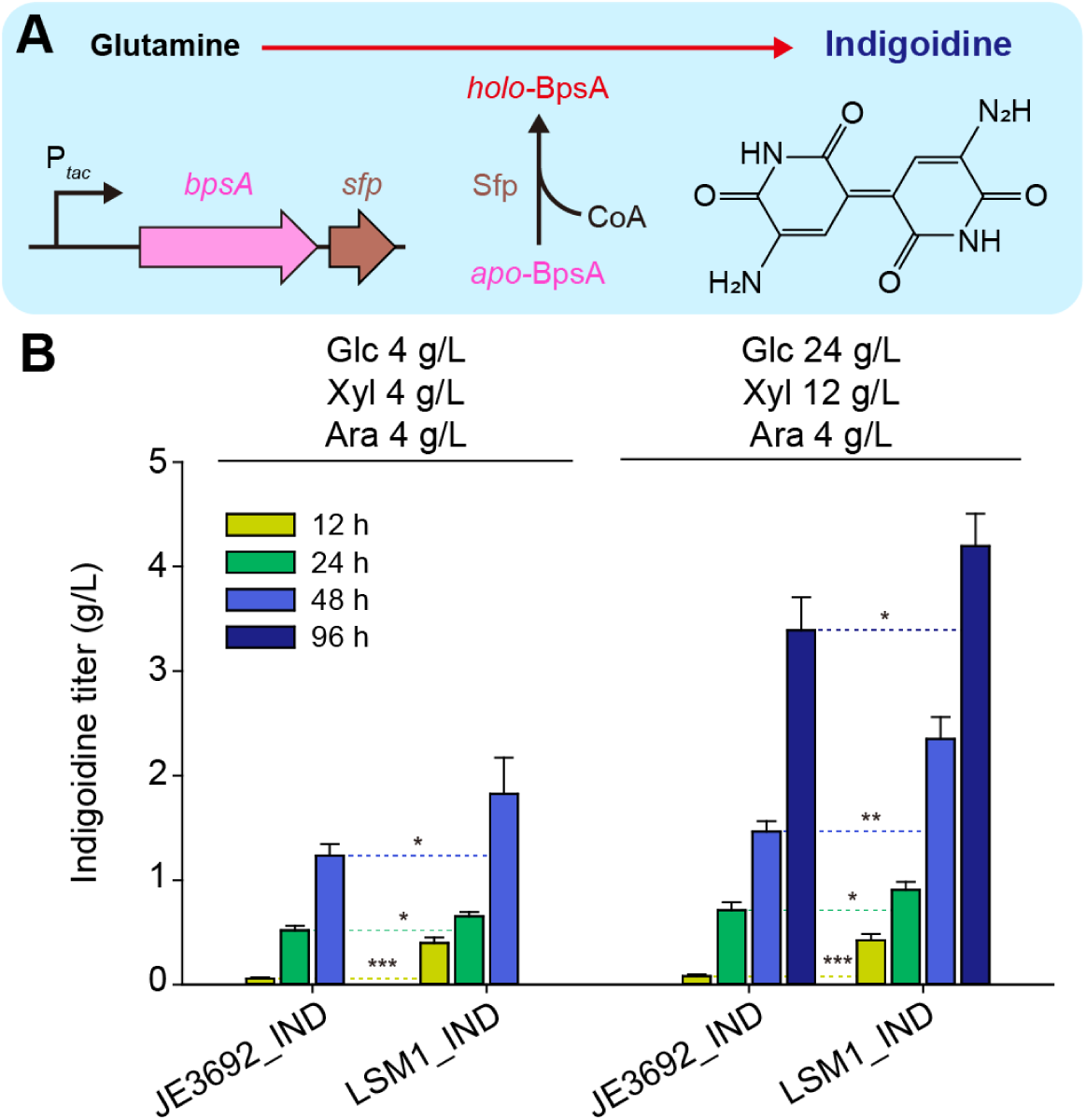
Indigoidine production from lignocellulosic sugars using evolved strain as a host. **(A)** The indigoidine production pathway in engineered *P. putida* strains. Indigoidine can be produced from glutamine by heterologous expression of *bpsA* from *S. lavendulae* and *sfp* from *B. subtilis*. These genes were expressed under the Tac promoter (P_­_) and the cassette was introduced into a plasmid. **(B)** Comparison of the indigoidine titers (g/L) of engineered strains based on the parental strain (JE3692) and an evolved strain (LSM1) after 12 h (yellow), 24 h (green), 48 h (blue), and 96 h (dark blue) cultivation under isotonic sugar condition (4 g/L of each sugar) and mock hydrolysate condition (24 g/L of glucose, 12 g/L of xylose, and 4 g/L of arabinose). The *x*- and *y*-axes indicate producer strain and indigoidine titer (g/L). *, ** and *** indicate statistical significance (*p* < 0.05, *p* < 0.01 and *p* < 0.001, respectively) between cultures of two different strains measured at the same timepoint. The cultures were conducted with biological triplicates (*n* = 3) and error bars indicate the standard deviations.

The LSM1_IND strain demonstrated a consistently higher indigoidine titer compared to the JE3692_IND strain (**Figure 5B**). Under isotonic conditions, the LSM1_IND strain produced 1.8 g/L of indigoidine over 48 hours, 1.5-fold higher compared to the JE3692_IND strain under the same conditions. When grown on mock hydrolysate, the LSM1_IND strain achieved a titer of 4.2 g/L over 96 hours, 1.2-fold higher compared to the JE3692_IND strain. This titer improvement was likely due to increased xylose and arabinose catabolism and metabolic flux redistribution described in the previous sections. Specifically, the efficient and direct supply of 2-ketoglutarate, resulting from enhanced arabinose catabolism, likely played a crucial role in increasing indigoidine production. The difference in titer between the two strains was more pronounced in the early stages of cultivation, with the LSM1_IND strain exhibiting 6.9-fold and 5.2-fold higher titers than the JE3692_IND strain after 12 hours under isotonic and mock hydrolysate conditions, respectively. These results demonstrate the utility of dynamic ALE for improving the conversion of mixed substrates into a single value-added product.

## Discussion

In this study, we conducted an automated ALE campaign on a previously engineered *P. putida* KT2440 strain, JE3692^23^, under various dynamic selection regimes to optimize lignocellulosic sugar catabolism. Statically evolved isolates generally exhibited improved growth relative to the starting strain only under their respective selection conditions. Among dynamically evolved isolates, those from the HSM and GXA regimes likewise evolved into catabolic specialists. In contrast, LSM isolates, evolved under conditions that required complete consumption of all three sugars, functioned as catabolic generalists with robust growth across single- and mixed-sugar environments (**Figure 2B, E**). These findings indicate that LSM is an effective dynamic selection regime for the simultaneous optimization of multi-sugar catabolism within a single ALE experiment and provide a useful framework for developing robust microbial hosts for mixed-carbon feedstock environments. Furthermore, the ALE-derived LSM1 strain led to improved production of indigoidine demonstrating utility for biomanufacturing goals (**Figure 5B**).

The distinct mutational landscapes and variations in catabolic performance across dynamic ALE regimes provided valuable insights into the rational design of evolutionary engineering strategies. Interestingly, despite being evolved under the mixed-sugar conditions, the HSM isolates displayed lower fitness than the LSM isolates when tested under the mixed-sugar conditions (**Figure 3D–E**). This outcome likely reflects differences in sugar concentrations and transfer criteria during the ALE experiments. Under the HSM mixed-sugar conditions, carbon remained abundant until transfer, likely favoring early sweeps of sprinters that rapidly dominated the population by utilizing easily catabolizable carbon sources^65^. However, such dynamics might not enforce sustained selection for a high overall growth rate across the desired time frame for applications. Consequently, HSM isolates often lost xylose catabolic ability that imposed fitness costs and evolved as specialists on glucose and arabinose. These results provide a mechanistic example of how fitness costs associated with specific catabolic pathways may contribute to hierarchical sugar utilization in nutrient-rich and stable environments^10,11^.

By contrast, LSM maintained low carbon availability and required complete sugar depletion before transfer, imposing selection for efficient use of all three sugars and promoting generalists^65^. Furthermore, disruptive mutations in flagellar biosynthetic genes (*fliR* and *flgH*) and associated regulatory genes (*fliA* and PP_5263 annotated as a GGDEF domain protein) in LSM isolates (**Supplementary Data 1**) are likely attributed to more efficient proteome reallocation and reduced motility or biofilm formation that increase specific growth rate^66,67^. Overall, these findings emphasize the need to carefully and rationally design ALE conditions that preserve and co-optimize targeted metabolic pathways while also accounting for factors such as mutation rates and population sizes^39^.

The investigation of selective disruption of the xylose catabolic pathway provided key insights into metabolic pathway optimization and evolutionary engineering outcomes. Evolutionary outcomes for the xylose pathway differed markedly between selective vs. non-selective regimes, reflecting a balance between maintaining catabolic function and minimizing resource expenditure. Under selective regimes, evolution appeared to prioritize reducing energy and resource burden over preserving maximal catabolic activity. Conversely, under non-selective regimes, strains frequently evolved with disruptions in the xylose pathway, likely due to the lack of selective pressure to maintain catabolic activity and the fitness cost imposed by constitutively overexpressed XylA and XylB. Consistent with our finding, several previous studies have reported that XylB overexpression impairs cell fitness in *Saccharomyces cerevisiae* and *Corynebacterium glutamicum*^68–70^. These divergent evolutionary outcomes highlight that evolutionary stability arises from the combined effects of pathway design, enzyme specificity, functional efficiency, and expression levels, which are especially important when optimizing multiple pathways.

This study represents a highly complex and parallelized ALE investigation, enabled by the use of automated robotic ALEbot platforms. Prior plate-based automated ALE studies typically relied on fixed schedules, such as transferring once per day or upon all cultures reaching a defined state, which limited diverse selection regimes, particularly under regimes such as LSM and HSM^71,72^. In this study, each lineage was cultivated in a separate tube, enabling individual OD tracking and timely transfers that maintained consistent selective regimes and promoted distinct evolutionary trajectories. Our results demonstrate that the dynamic ALE regime that enforced complete consumption of all sugars can optimize multi-substrate catabolism, highlighting their advantage over traditional static ALE that relies on a single selection pressure. Looking ahead, we expected that this ALE-based dynamic selection strategy could be systematically applied to even more complex and heterogeneous feedstocks, such as lignin-derived aromatics, marine polysaccharides, and depolymerized mixed plastic waste^43,73–77^.

## Methods

### Bacterial strains, plasmids, and reagents

Strains and plasmids used in this study are listed in **Supplementary Table 2**. Plasmids were primarily constructed using the NEBuilder HiFi DNA Assembly Kit (New England Biolabs, Ipswich, MA, USA). Oligonucleotides, synthesized by Integrated DNA Technologies (IDT, Coralville, IA, USA), are listed in **Supplementary Table 3**. The plasmids used for recombination were based on the pK18mobsacB backbone (ATCC 87097) and recombination regions either cloned from isolates ARA3, LSM1, LSM2 or synthesized by GenScript (Piscataway, NJ, USA). For DNA amplification, the Q5 High-Fidelity DNA polymerase or OneTaq DNA polymerase (NEB) was used. Plasmids were purified using a Zymo PURE Plasmid Miniprep kit from Zymo Research (Irvine, CA, USA). Genomic DNA samples were prepared by using a Quick-DNA Fungal/Bacterial Miniprep kit from Zymo Research. Genome sequencing library samples were prepared using an NEBNext Ultra II kit from NEB (Ipswich, MA, USA). All chemicals were purchased from Sigma-Aldrich (St. Louis, MO, USA) and used without further purification.

### Culture conditions

Bacterial cultivations were conducted in a Luria−Bertani (LB, 10 g/L tryptone, 5 g/L yeast extract, 10 g/L NaCl) or modified minimal M9 medium (4 g/L glucose, 2 g/L (NH_­_)_­_SO_­_, 6.8 g/L Na_­_HPO_­_, 3 g/L KH_­_PO_­_, 0.5 g/L NaCl, 2 mM MgSO_­_, 0.1 mM CaCl_­_, 500 μL/L 2,000× trace element solution). The composition of the trace element solution is 4.5 g/L ZnSO_­_·7H_­_O, 0.7 g/L MnCl_­_·4H_­_O, 0.3 g/L CoCl_­_·6H_­_O, 0.2 g/L CuSO_­_·2H_­_O, 0.4 g/L Na_­_MoO_­_·2H_­_O, 4.5 g/L CaCl_­_·2H_­_O, 3.0 g/L FeSO_­_·7H_­_O, 1.0 g/L H_­_BO_­_, 0.1 g/L KI, 15 g/L disodium ethylenediaminetetraacetate^42,78^. For production strains only, gentamicin was supplemented at 30 μg/mL to maintain the plasmid carrying the indigoidine biosynthetic genes.

Cultures were performed in 30 mL cylindrical tubes containing 15 mL of medium at 30 °C, agitated at 1,100 rpm with a magnetic stirring bar. Seed cultures were prepared by inoculating colonies from LB agar plates into the minimal medium and incubating them overnight. Grown cultures were diluted into the fresh medium at an OD_­_of 0.1. When OD_­_reached 0.6 ∼ 1, cells were harvested and re-inoculated into minimal media at OD_­_of 0.05 to initiate experimental cultures.

For ALE, 150 μL of cultures at the late-exponential phase (OD_­_of 0.6 ∼ 1) were passed into minimal medium supplemented with the indicated carbon sources, using an automated robotic platform^39^. OD_­_was measured using a Sunrise™ microplate reader from Tecan (Männedorf, Switzerland). A maximum specific growth rate (h^-1^) was calculated by linear regression between ln(OD_­_) and time (hour) during the exponential growth phase (OD_­_∼ 1).

### Genome resequencing

Raw sequencing reads were obtained by using a NovaSeq 6000 (Illumina) at the UC San Diego IGM Genomics Center and analyzed by using Breseq (version 0.35.4)^79^ and Bowtie2 (version 2.2.6)^80^. Mutations consistently present in the starting strain were filtered out (**Supplementary Table 4**). Analysis results were uploaded to ALEdb v1.0^81^ and are accessible at http://aledb.org (project id: Pputida_dynamic_ALE). The summarized analysis results are also included in **Supplementary Data 1**. Raw genome-sequencing files are available at the NCBI SRA database (BioProject number: PRJNA1395279, **Supplementary Table 5**).

### Protein expression and purification

Genes encoding XylA and XylB were cloned into a pET15b vector and transformed into *E. coli* BL21(DE3). Cultures were grown at 37 °C in LB medium containing 50 µg/mL ampicillin until reaching an OD_­_of 0.6–0.8, at which point expression was induced with 1 mM IPTG and continued overnight at 18 °C. Cells were harvested by centrifugation at 4,000 × *g* for 30 min at 4 °C, resuspended in lysis buffer (20 mM TRIS pH 8, 300 mM NaCl, 5 mM imidazole) and lysed by sonication on ice. Lysates were clarified by centrifugation and purified using nickel affinity chromatography (Ni-NTA IMAC). Eluted fractions were dialyzed (20 mM TRIS pH 8, 300 mM NaCl), concentrated to >10 mg/mL, and flash frozen dropwise in liquid nitrogen, then stored at - 80 °C until use. XylA concentration was determined by absorbance at 280 nm using a calculated extinction coefficient of 71,070 M^-1^cm^-1^, and XylB concentration was determined by absorbance at 280 nm using a calculated extinction coefficient of 82,756 M^-1^cm^-1^.

### NMR analysis of XylA activity

Reactions were performed in 100 mM potassium phosphate buffer (pH 8) containing 10 mM MgSO_­_, 20 mM sugar substrate (D-xylose, D-glucose, or L-arabinose), and 12 µM purified XylA. Reaction mixtures (final volume: 600 µL) were supplemented to reach 10% D_­_O and 0.1 mg/mL TMSP (trimethylsilyl propionic acid) as an internal chemical shift and integration reference. ^1^H NMR spectra were acquired on a 400 MHz NMR spectrometer at room temperature using a standard one-dimensional water-suppression pulse sequence. Spectra were processed and analyzed in Mestrenova (Mestrelab Research). Product peaks for D-xylulose and D-fructose were identified based on available published or reference spectra and quantified by peak integration. The L-ribulose product lacked published assignments; thus, quantification was based on a representative signal assumed to correspond to one or more protons (≥1 H). Final concentrations were estimated using relative integration values calibrated to the TMSP internal standard.

### Coupled enzymatic assay to detect XylB activity

XylB activity was monitored using a continuous, coupled assay with pyruvate kinase (PK) and lactate dehydrogenase (LDH) linking ATP consumption to NADH oxidation. Reactions were conducted in 200 mM potassium phosphate buffer (pH 8.0) supplemented with 20 mM MgSO_­_, 2 mM phosphoenolpyruvate (PEP), 0.01X PK/LDH enzyme mix (with the 1X mixture corresponding to ∼600–1,000 U/mL pyruvate kinase and ∼900–1400 U/mL LDH), 1 mM NADH, and 0.04 µM purified XylB. Sugar substrates were added at 10 mM final concentration, except D-xylulose, which was added at 2 mM. The final reaction volume was 200 µL and reactions were performed in 96-well UV-STAR microplates (Greiner Bio-One). Absorbance at 340 nm (A_­_) was monitored at 1-minute intervals for 60 minutes at room temperature using a BioTek Synergy H1 plate reader.

### Proteome analysis

*P. putida* cultures were collected after the OD_­_reached 0.6 in minimal media supplemented with 4 g/L of glucose. Cells were harvested and stored at -80 °C until further processing. Protein was extracted from cell pellets and tryptic peptides were prepared following the established proteomic sample preparation protocol^82^. Briefly, cell pellets were resuspended in Qiagen P2 Lysis Buffer (Qiagen, Germany) to promote cell lysis. Proteins were precipitated with the addition of 1 mM NaCl and 4 x vol acetone, followed by two additional washes with 80% acetone in water. The recovered protein pellet was homogenized by pipetting mixing with 100 mM ammonium bicarbonate in 20% methanol. Protein concentration was determined by the DC protein assay (BioRad, USA). Protein reduction was accomplished using 5 mM tris 2-(carboxyethyl)phosphine (TCEP) for 30 min at room temperature, and alkylation was performed with 10 mM iodoacetamide (IAM; final concentration) for 30 min at room temperature in the dark. Overnight digestion with trypsin was accomplished with a 1:50 trypsin:total protein ratio.

The resulting peptide samples were analyzed on an Agilent 1290 UHPLC system coupled to a Thermo Scientific Orbitrap Exploris 480 mass spectrometer for discovery proteomics^83^. Briefly, peptide samples were loaded onto an Ascentis® ES-C18 Column (Sigma–Aldrich, USA) and were eluted from the column by using a 10-minute gradient from 98% solvent A (0.1 % FA in H2O) and 2% solvent B (0.1% FA in ACN) to 65% solvent A and 35% solvent B. Eluting peptides were introduced to the mass spectrometer operating in positive-ion mode and were measured in data-independent acquisition (DIA) mode with a duty cycle of 3 survey scans from m/z 380 to m/z 985 and 45 MS2 scans with precursor isolation width of 13.5 m/z to cover the mass range.

DIA raw data files were analyzed by the integrated software suite DIA-NN^84^. The databases used in the DIA-NN search (library-free mode) are *P. putida* latest Uniprot proteome FASTA sequences plus the protein sequences of the heterologous proteins and common proteomic contaminants. DIA-NN determines mass tolerances automatically based on first pass analysis of the samples with automated determination of optimal mass accuracies. The retention time extraction window was determined individually for all MS runs analyzed via the automated optimization procedure implemented in DIA-NN. Protein inference was enabled, and the quantification strategy was set to Robust LC = High Accuracy. Output main DIA-NN reports were filtered with a global FDR = 0.01 on both the precursor level and protein group level. The Top3 method, which is the average MS signal response of the three most intense tryptic peptides of each identified protein, was used to plot the quantity of the targeted proteins in the samples^85,86^. The generated mass spectrometry proteomics data have been deposited to the ProteomeXchange Consortium via the PRIDE partner repository with the dataset identifier PXD055215^87^. The processed data and analysis results are provided in **Supplementary Data 2**. DIA-NN is freely available for download from https://github.com/vdemichev/DiaNN.

### Metabolite quantification

Indigoidine was quantified after its extraction using dimethyl sulfoxide as described previously^56^. Absorbance at 612 nm was measured by using an Infinite^®^ 200 PRO microplate reader from Tecan (Männedorf, Switzerland), and the concentration was determined from a calibration curve generated with indigoidine standards of known concentrations (**Supplementary Figure 9**). Concentrations of sugars were quantified by using a 1260 Infinity II LC system (Agilent, Santa Clara, CA, USA) equipped with an HPX-87H (Biorad, Hercules, CA, USA) and a refractive index detector. 5 mM H_­_SO_­_was used as a mobile phase at a flow rate of 0.5 mL/min. The column temperature was maintained at 60 °C. A sugar consumption rate (g/L/h) was calculated by dividing the amount of sugar consumed (g/L) up to the late exponential phase by the corresponding cultivation time (h).

## Acknowledgements

This material is based upon work at the Joint BioEnergy Institute supported by the U.S. Department of Energy, Office of Science, Biological and Environmental Research under Award Number DE-AC02-05CH11231. This work was also authored in part by the National Renewable Energy Laboratory for the U.S. Department of Energy (DOE) under Contract Number DE-AC36-08GO28308. Funding was provided to BNB, TML, CWJ, and GTB by the U.S. DOE Office of Energy Efficiency and Renewable Energy Bioenergy Technologies Office (BETO) Agile BioFoundry via Contract Number DE-AC36-08GO28308. This work was also authored in part by Oak Ridge National Laboratory, which is managed by UT-Battelle, LLC, for the U.S. Department of Energy (DOE) under Contract Number DE-AC05-00OR22725. Funding was provided to NG and AMG by the BETO Agile BioFoundry. The United States Government retains and the publisher, by accepting the article for publication, acknowledges that the United States Government retains a nonexclusive, paid-up, irrevocable, worldwide license to publish or reproduce the published form of this manuscript, or allow others to do so, for United States Government purposes. Any subjective views or opinions that might be expressed in this paper do not necessarily represent the views of the U.S. Department of Energy or the United States Government. H.G.L. acknowledges the National Research Foundation of Korea (NRF) grants funded by the Korea government (MSIT) (grant number RS-2024-00334792 and RS-2025-02215308). We thank Gayle Bentley and Lisa Guay at DOE and members of the Agile BioFoundry for helpful discussions. The purified indigoidine used in this study was kindly prepared by Javier Menasalvas. We thank Ciara de Venecia, Elina A. Olson, and Richard Szubin for technical support.

