## Supplementary Information for "Simultaneous optimization of lignocellulosic sugar catabolism via systematic laboratory evolution in dynamic conditions"

**Contents**

Number of pages: 17

Number of figures: 9

Number of tables: 5

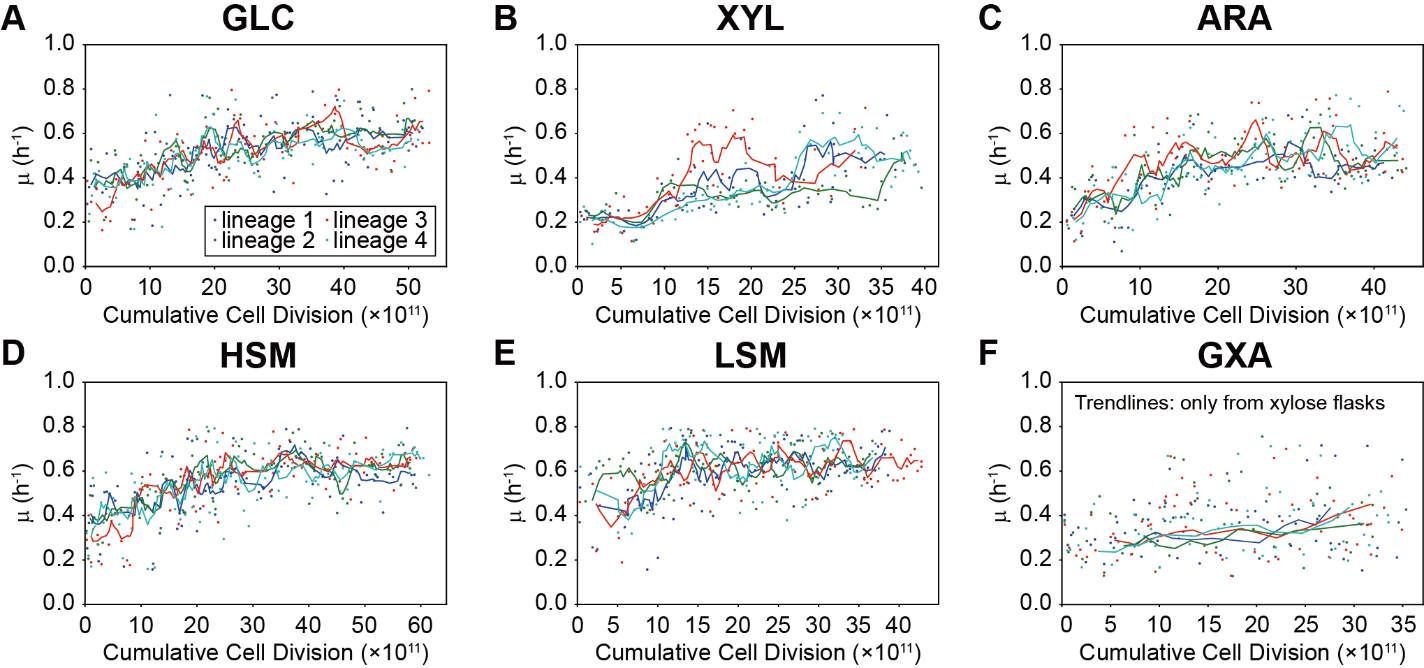

**Supplementary Figure 1. ALE trajectories of statically and dynamically evolved lineages**

Growth trajectories during the ALE under (A) GLC, (B) XYL, (C) ARA, (D) HSM, (E) LSM, and (F) GXA selection regimes. Scatter plots indicate the specific growth rate observed at each passage according to cumulative cell division (CCD). Line plots represent the moving average trendline for every 5 points. For GXA, only the trendline from the xylose flasks, which showed the only increasing trend, was depicted. The *x*- and *y*-axes indicate CCD and specific growth rate (h^-1^), respectively. Each color represents replicated ALE experiments.

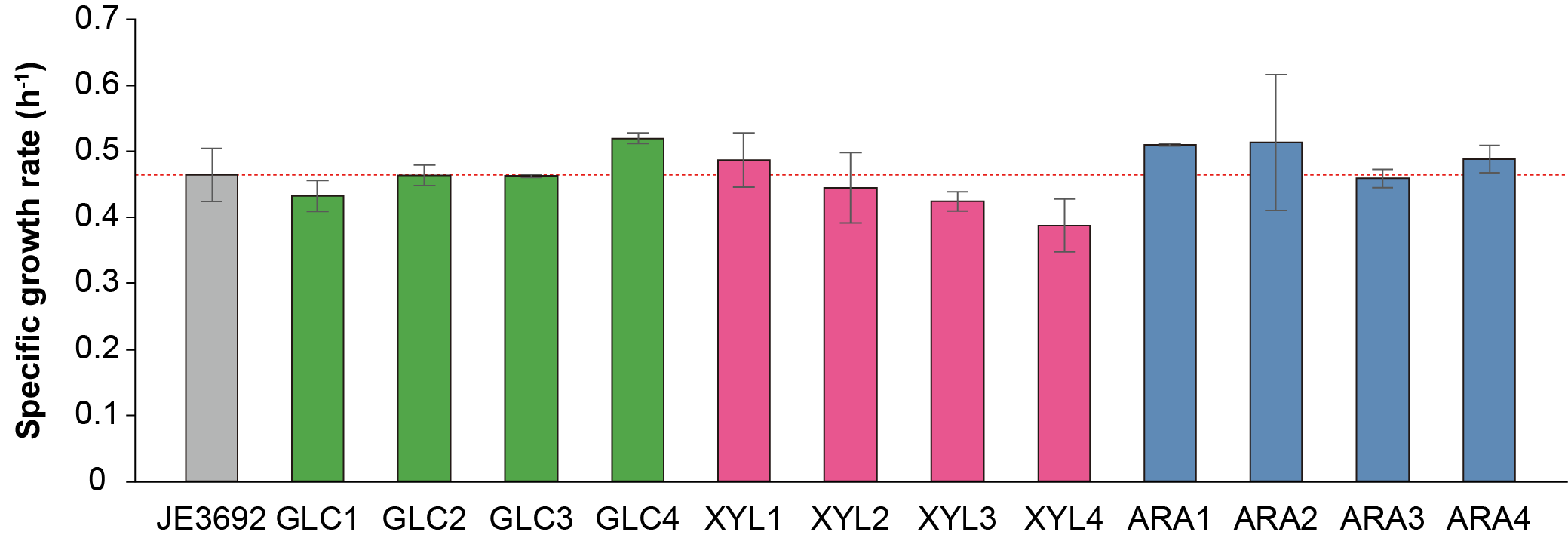

**Supplementary Figure 2. Specific growth rates of statically evolved strains under the HSM mixed-sugar conditions.**

The *x*- and *y*-axes indicate strains and specific growth rate (h^-1^), respectively. The gray, green, pink, and boxes represent JE3692, GLC, XYL, and ARA isolates, respectively. Red dotted lines indicate the growth rate of the JE3692 strain. The cultures were conducted with three biological replicates (*n* = 3), and error bars indicate the standard deviations.

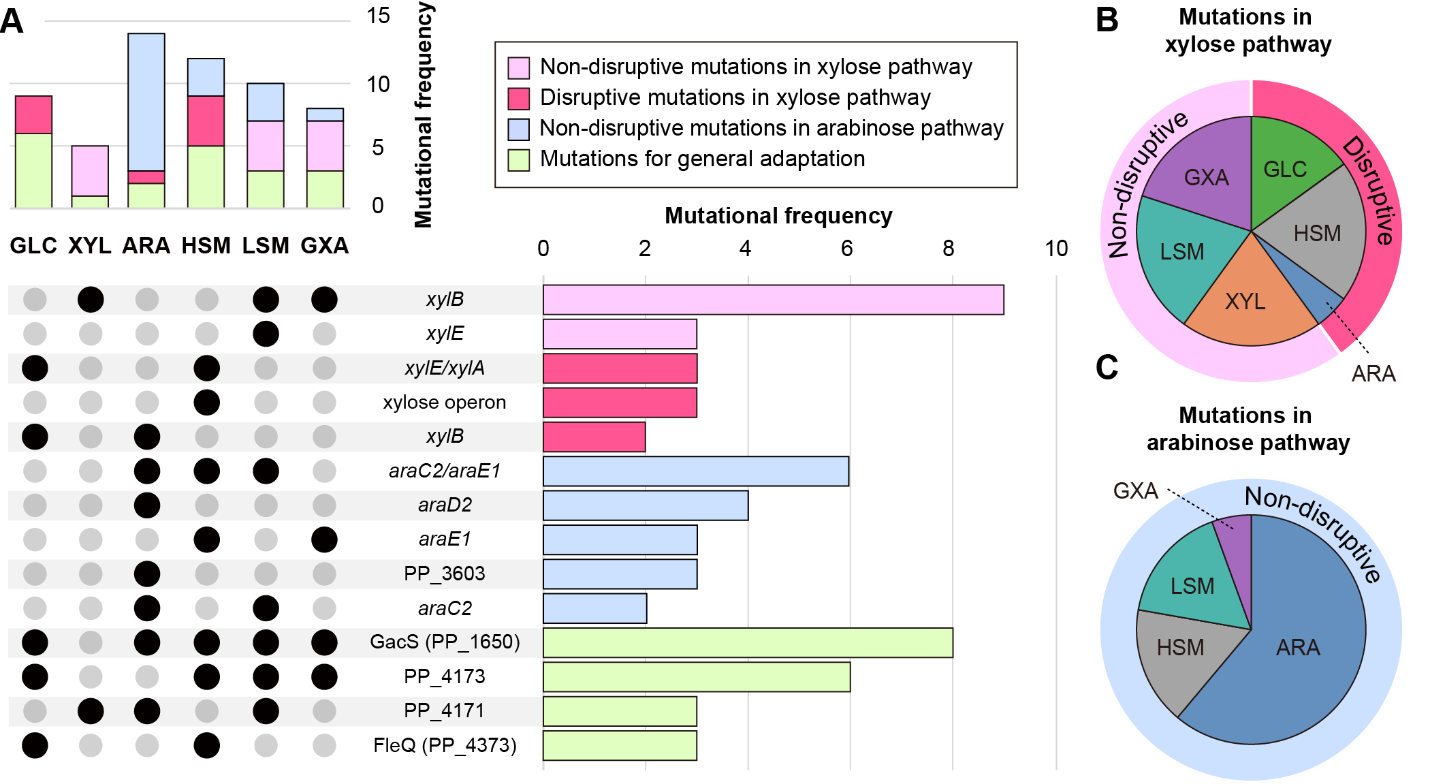

**Supplementary Figure 3. Distribution of mutational frequencies across different ALE groups**

(A) UpSet plots to summarize the mutational patterns across the ALE groups. The top vertical bar chart represents the mutational frequency by ALE group. The bottom right horizontal bar chart represents the mutational frequency of the sets sharing mutational genes or regions, which are specified by black dots in the bottom left. Pie plots showing the mutational frequencies identified in the (B) xylose and (C) arabinose pathways across each ALE group and their impact on functionality. The color code of the outer pie plots in panel B and C is the same as in the bar plots in panel A.

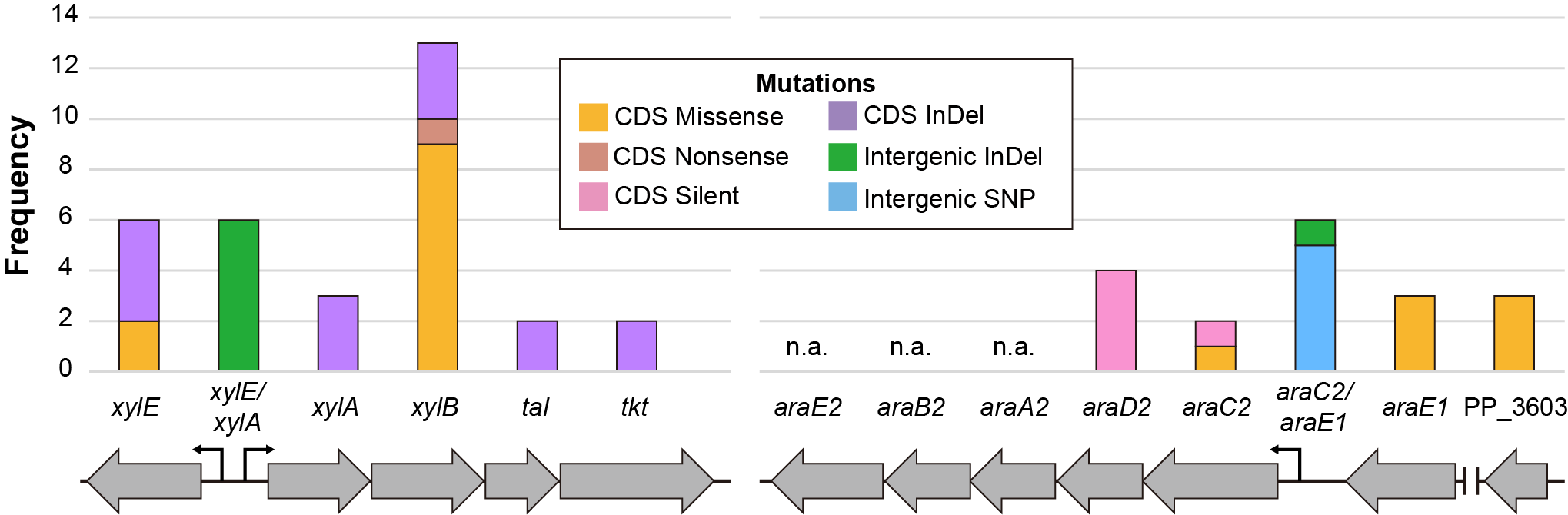

**Supplementary Figure 4. Mutational frequency distribution across genetic regions of the xylose and arabinose catabolic pathways**

Frequency of mutational types in regions related to catabolism of xylose and arabinose. Different colors represent different mutation types: orange, single nucleotide variants (SNVs) in CDS causing missense mutations; brown, SNVs in CDS causing nonsense mutations; pink, SNVs in CDS causing silent mutations; purple, insertion or deletion mutations (InDels) in CDS; light blue, SNVs in intergenic region; green, InDels in intergenic regions. The *x*- and *y*-axes indicate genetic region and mutational frequency. The arrows below the *x*-axis depict the genetic context.

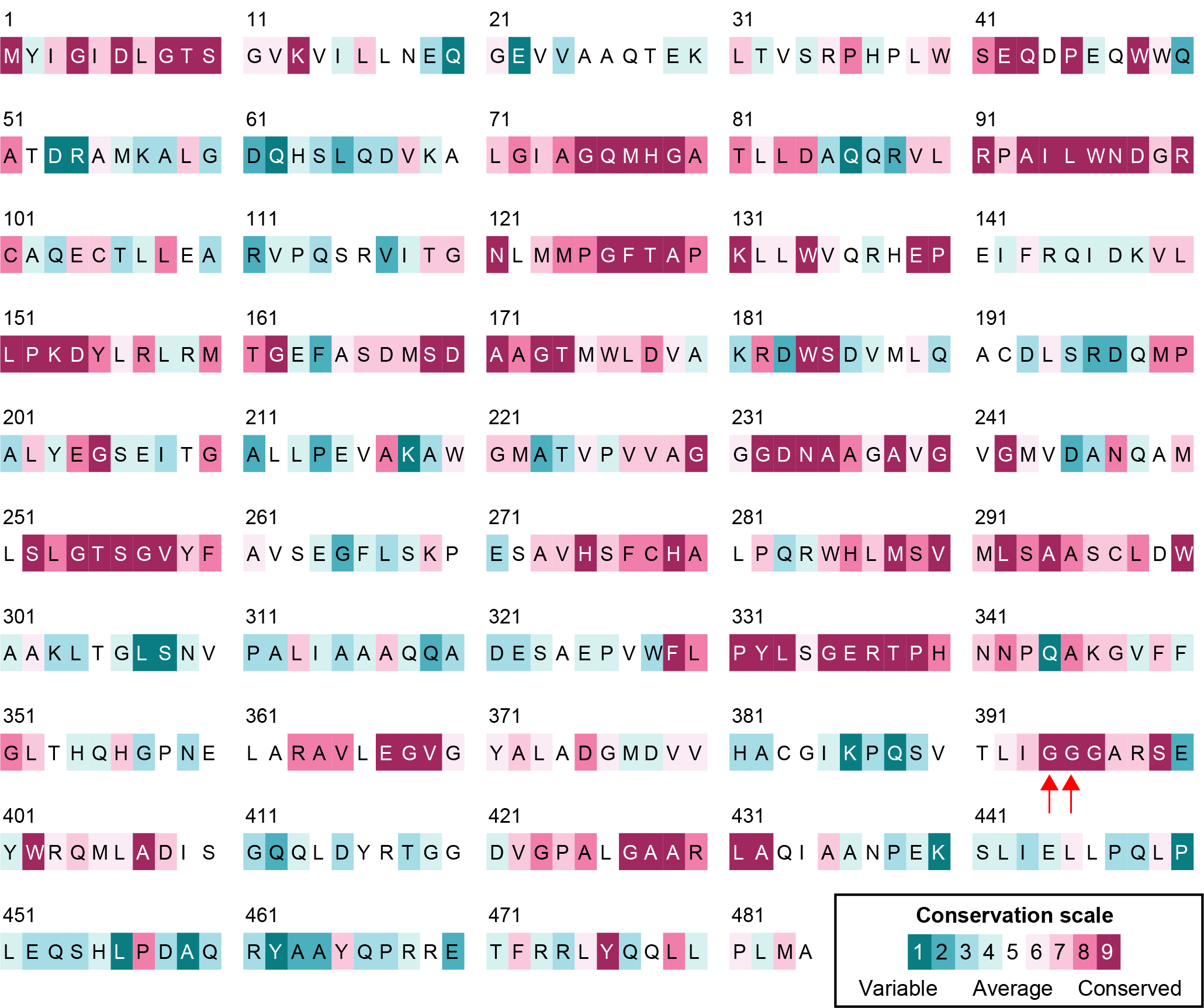

**Supplementary Figure 5. Conservation analysis of XylB**

The degree of conservation of amino acids was shown in the coloring scheme. The color intensity increases based on amino acids conservation grades e.g. turquoise indicates variable sites; white indicates average sites; maroon indicates evolutionarily conserved sites. The convergently mutated residues (395 and 396) are indicated by red arrows.

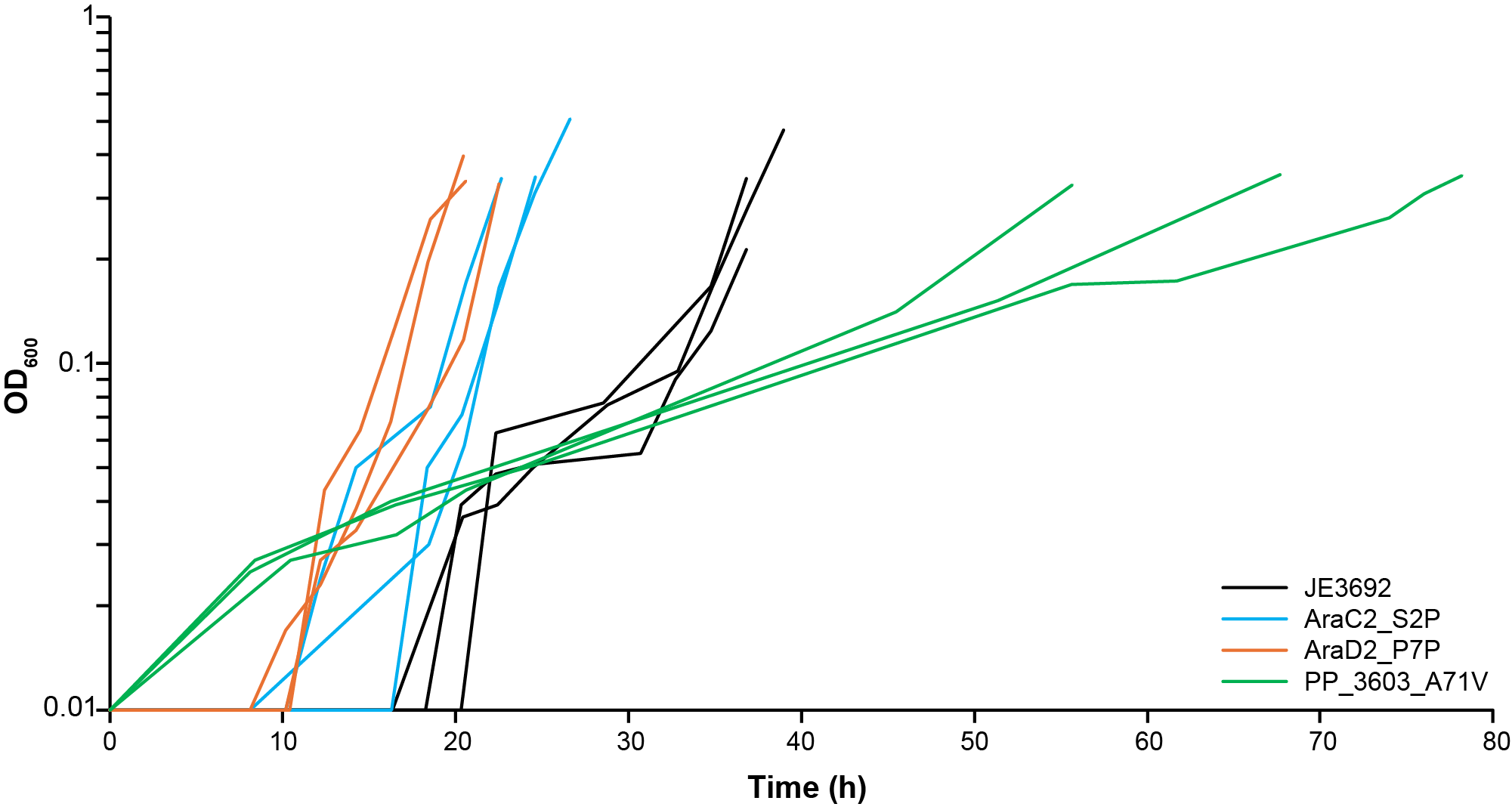

**Supplementary Figure 6. Differences of growth profiles under arabinose condition by presence of key mutations.**

Growth profiles of the JE3692 (black), AraC2_S2P (blue), AraD2_P7P (orange), and PP_3603_A71V (green) strains in minimal medium supplemented with 4 g/L arabinose. The x- and *y*- axis indicate time (h) and cell biomass (OD_600_), respectively. The cultures were conducted with three biological replicates (*n* = 3).

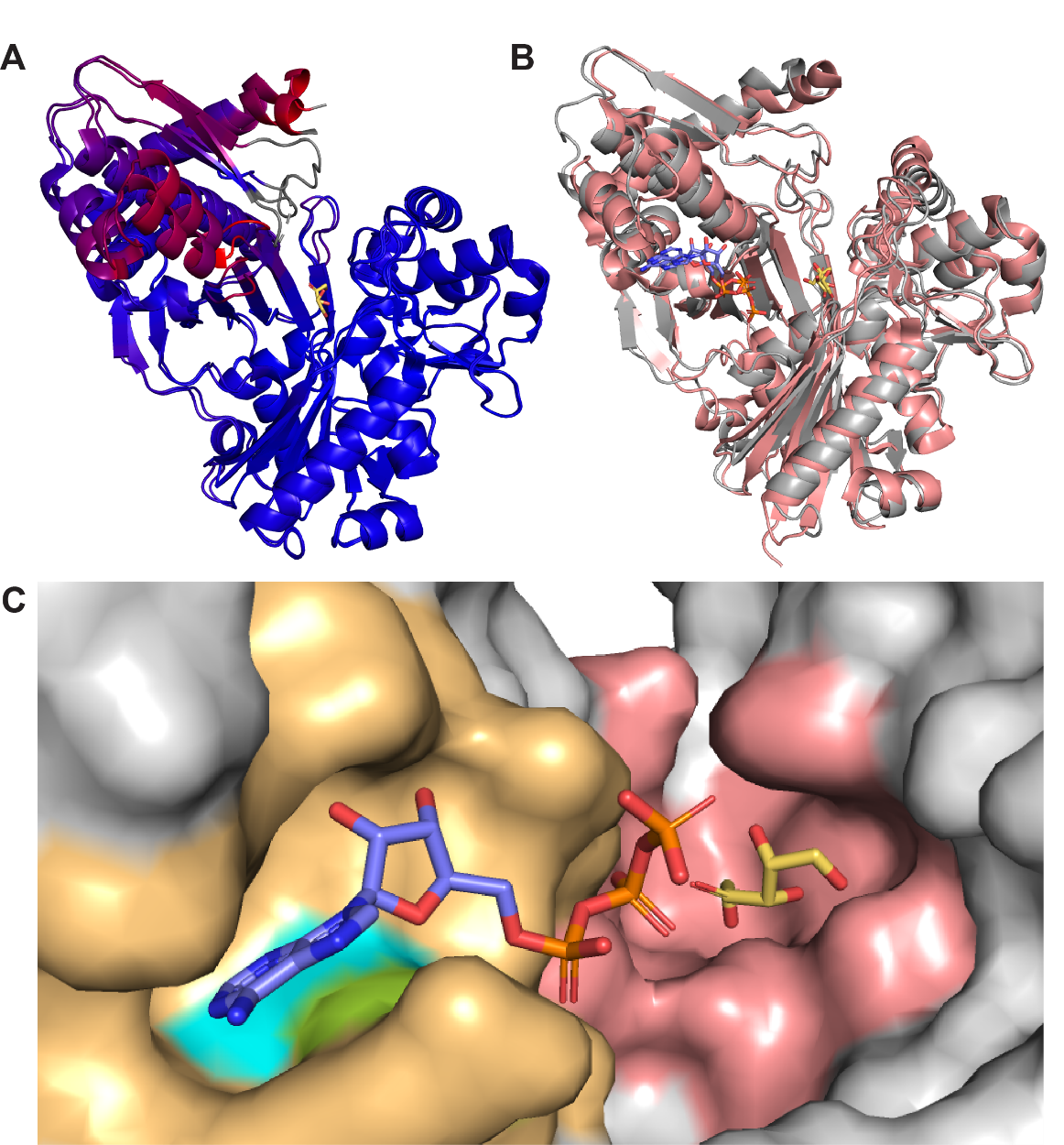

**Supplementary Figure 7. Generation of an Alphafold 3 model of *E. coli* XylB with ATP and D-xylulose.**

(A) Alignment of the E. coli XylB crystal structure (PDB: 2ITM) complexed with D-xylulose (XUL) with the AlphaFold3 (AF3) model (all-atom RMSD = 0.625 Å), showing close alignment at the N-terminal region and the XUL binding site. This structural correspondence supports transferring the XUL ligand from the 2ITM crystal structure to the AF3 model. Structures are colored by RMSD, with blue indicating lower deviation and red indicating higher deviation. (B) Alignment of a XylB homolog crystal structure from *Lactobacillus acidophilus* (PDB: 3LL3) complexed with ATP and XUL with the Alphafold3 structure of E. coli XylB with ATP docked (all-atom RMSD = 1.573 Å). The conserved placement of ATP in 3LL3 supports the ATP docking generated by the AF3 webserver. (C) AF3 model of E. coli XylB with ATP (positioned by AF3 docking) and XUL (transferred from the 2ITM structure). Residues within 6 Å are highlighted for ATP (orange) and XUL (red). The mutated residues G395 and G396, shown in cyan and green, respectively, are located in the ATP binding site. For visual clarity of all the displayed XylB structures, only one monomer of the dimeric biological assembly is shown.

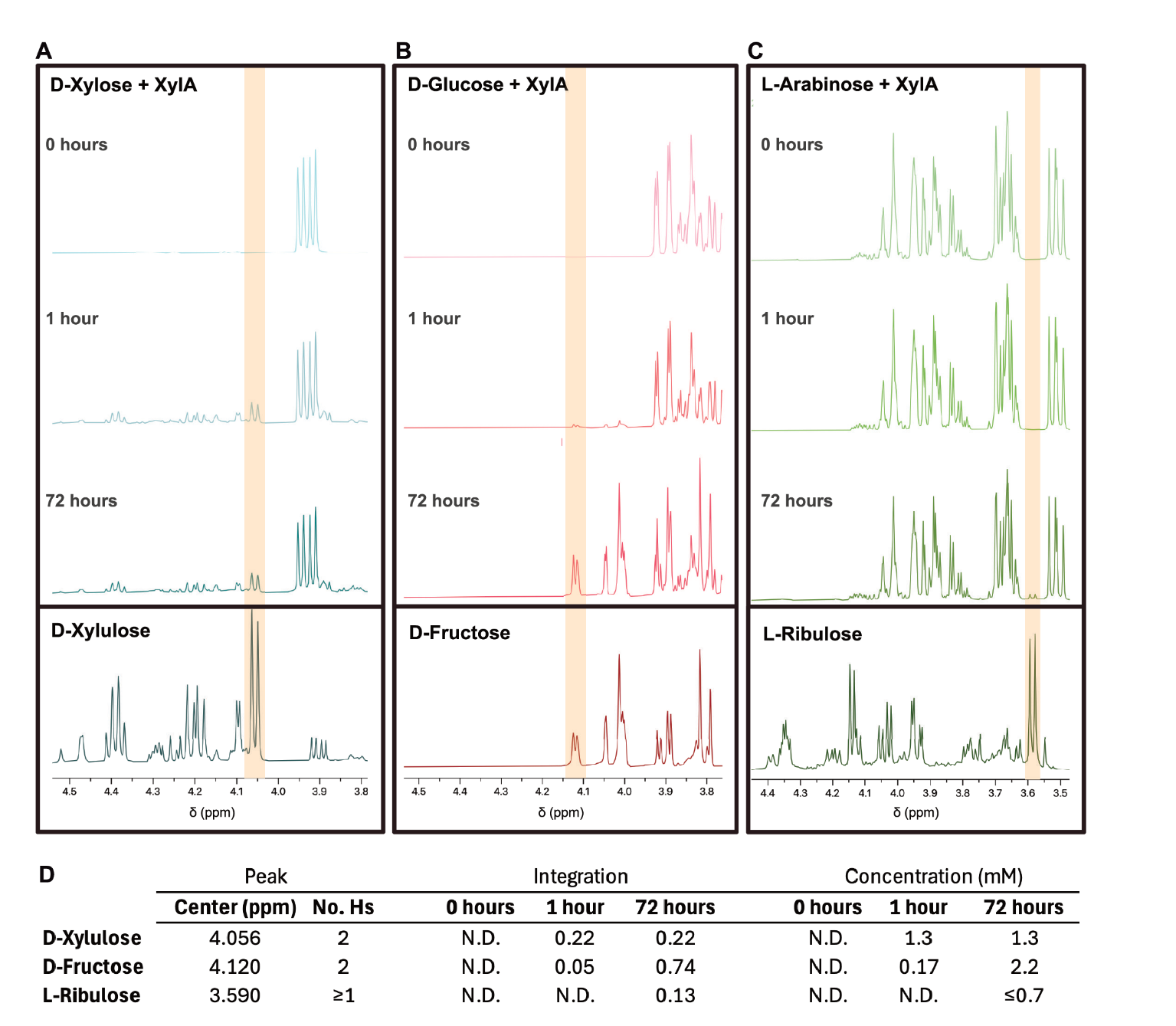

**Supplementary Figure 8. NMR analysis of XylA activity on various sugar substrates**

Time-course proton NMR spectra of reactions containing recombinant XylA and either (A) D-xylose, (B) D-glucose, or (C) L-arabinose over 72 hours. Each panel includes spectra at 0, 1, and 72 hours after XylA addition. Spectra for D-xylulose, D-fructose, and L-ribulose standards are shown for comparison. The shaded regions highlight a unique product peak for quantification against an internal standard. (D) Summary of representative product peaks including chemical shift, number of hydrogens, signal integration, and calculated concentrations based on comparison to an internal standard at 1 hour and 72 hours. D-xylulose formation from D-xylose is observed rapidly and remains stable. D-fructose accumulates gradually in D-glucose reactions. Minor L-ribulose formation is detected from L-arabinose. “N.D.” indicates signal not detected above noise. No conversion was observed for samples without XylA (data not shown).

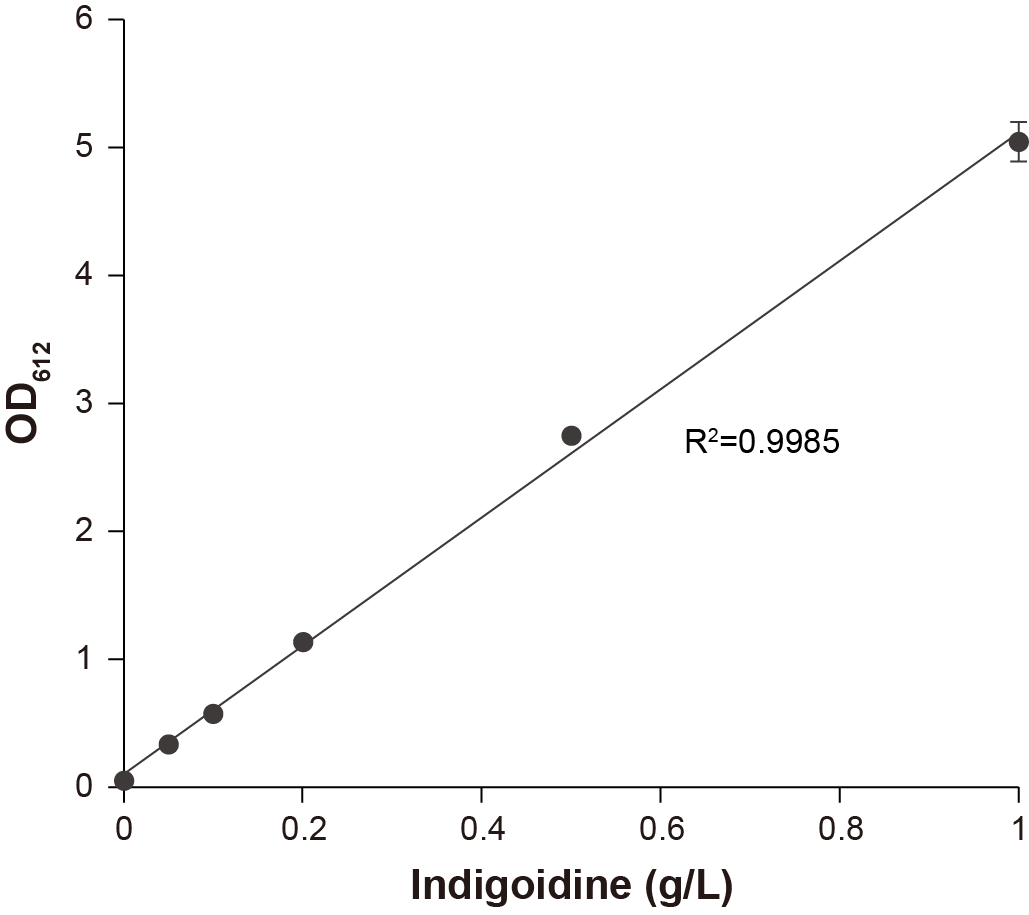

**Supplementary Figure 9. Standard curve for indigoidine quantification**

Indigoidine was dissolved at various concentrations in the minimal medium and the absorbance at 612 nm was measured. Error bars indicate standard deviations of three technical replicates.

**Supplementary Table 1. Summary of the ALE experiment results**

| **ALE group** | **Selection regime** | **Initial growth rate (h^-1^)** | **Replicate** | **# of  passages** | **# of generations** | **Cumulative  cell divisions  (CCD, 10^11^)** | **Growth rate of  the end-point  population (h^-1^)** |
| --- | --- | --- | --- | --- | --- | --- | --- |
| GLC | Glucose 4 g/L | 0.42 ± 0.04 | 1 | 97 | 634 | 52.2 | 0.62 |
|  |  |  | 2 | 98 | 639 | 52.0 | 0.63 |
|  |  |  | 3 | 97 | 631 | 53.2 | 0.56 |
|  |  |  | 4 | 95 | 618 | 51.5 | 0.57 |
| XYL | Xylose 4 g/L | 0.20 ± 0.01 | 1 | 65 | 394 | 36.6 | 0.44 |
|  |  |  | 2 | 67 | 413 | 39.6 | 0.36 |
|  |  |  | 3 | 70 | 417 | 33.1 | 0.44 |
|  |  |  | 4 | 69 | 413 | 39.3 | 0.43 |
| ARA | Arabinose 4 g/L | 0.14 ± 0.02 | 1 | 74 | 479 | 42.3 | 0.40 |
|  |  |  | 2 | 78 | 491 | 44.1 | 0.44 |
|  |  |  | 3 | 81 | 519 | 43.7 | 0.48 |
|  |  |  | 4 | 80 | 514 | 43.8 | 0.51 |
| HSM | Glucose 4 g/L+ Xylose 4 g/L+ Arabinose 4 g/L (Selective consumption allowed) | 0.46 ± 0.04 | 1 | 105 | 687 | 58.9 | 0.58 |
|  |  |  | 2 | 106 | 692 | 59.1 | 0.52 |
|  |  |  | 3 | 107 | 692 | 59.2 | 0.66 |
|  |  |  | 4 | 109 | 709 | 61.5 | 0.58 |
| LSM | Glucose 0.5 g/L+ Xylose 0.5 g/L+ Arabinose 0.5 g/L (Complete consumption mandated) | 0.46 ± 0.04 | 1 | 103 | 671 | 38.8 | 0.58 |
|  |  |  | 2 | 96 | 627 | 37.1 | 0.68 |
|  |  |  | 3 | 106 | 689 | 42.8 | 0.62 |
|  |  |  | 4 | 95 | 617 | 37.5 | 0.61 |
| GXA | Glucose 4 g/L/ Xylose 4 g/L/ Arabinose 4 g/L (Alternating supply) | Glucose: 0.42 ± 0.04 Xylose: 0.20 ± 0.01 Arabinose: 0.14 ± 0.02 | 1 | 63 | 409 | 32.4 | Glucose: 0.67 Xylose: 0.25 Arabinose: 0.15 |
|  |  |  | 2 | 70 | 447 | 35.4 | Glucose: 0.65 Xylose: 0.37 Arabinose: 0.21 |
|  |  |  | 3 | 68 | 441 | 34.9 | Glucose: 0.57 Xylose: 0.44 Arabinose: 0.22 |
|  |  |  | 4 | 68 | 437 | 33.5 | Glucose: 0.51 Xylose: 0.40 Arabinose: 0.27 |

**Supplementary Table 2. List of strains and plasmids used in this study**

| **Name** | **Description** | **Source** |
| --- | --- | --- |
| **Strains** |  |  |
| JE212 | *P. putida* KT2440 ∆*hsdR::Bxb1int-attB* ∆*gcd* | 1 |
| JE3226 | *P. putida* KT2440 ∆*hsdR::Bxb1int-attB* ∆*gcd* ∆*ampC::PxylE*-xylE:Ptac-xylAB:talB:tktA* | 1 |
| JE3285 | *P. putida* KT2440 ∆*hsdR::Bxb1int-attB* ∆*gcd::PxylE-araE1:Ptac-araC2D2A2B2E2* ∆*ampC::PxylE*-xylE:Ptac-xylAB:talB:tktA* | 1 |
| JE3692 | *P. putida* KT2440 *∆hsdR::Bxb1int-attB ∆gcd::PxylE-araE1:Ptac-araC2D2A2B2E2 ∆ampC::PxylE*-xylE:Ptac-xylAB:talB:tktA ∆crc* | 1 |
| BL21(DE3) | *E. coli* expression host for XylA and XylB | NEB |
| GLC1-4 | Isolates from evolved lineages under GLC ALE regime | This study |
| XYL1-4 | Isolates from evolved lineages under XYL ALE regime | This study |
| ARA1-4 | Isolates from evolved lineages under ARA ALE regime | This study |
| HSM1-4 | Isolates from evolved lineages under HSM ALE regime | This study |
| LSM1-4 | Isolates from evolved lineages under LSM ALE regime | This study |
| GXA1-4 | Isolates from evolved lineages under GXA ALE regime | This study |
| XylB_G395W | JE3692 *xylB* p.Gly395Trp | This study |
| XylE_Y179E | JE3692 *xylE* p.Tyr179Glu | This study |
| AraC2_S2P | JE3692 *AraC2* p.Ser2Pro | This study |
| AraD2_P7P | JE3692 *AraD2* p.Pro7Pro | This study |
| PP_3603_A71V | JE3692 PP_3603 p.Ala71Ala | This study |
| JE3692_IND | JE3692/pTE252_IND | This study |
| LSM1_IND | LSM1/pTE252_IND | This study |
| **Plasmids** |  |  |
| pET15b | Expression vector for XylA and XylB | Novagen |
| pEUK238 | pET15b_XylA | This study |
| pEUK239 | pET15b_XylB | This study |
| pEUK239_1 | pET15b_XylB_G395R | This study |
| pEUK239_2 | pET15b_XylB_G395W | This study |
| pNEG0020 | Recombination plasmid XylB_G395W, based on pK18mobsacB | This study |
| pNEG0024 | Recombination plasmid XylE_Y179C, based on pK18mobsacB | This study |
| pNEG0025 | Recombination plasmid AraC_S2P, based on pK18mobsacB | This study |
| pNEG0021 | Recombination plasmid AraD_P7P, based on pK18mobsacB | This study |
| pNEG0023 | Recombination plasmid PP_3603_A71V, based on pK18mobsacB | This study |
| pTE251 | pBBR1_*oriV*_*nptII*_*araC_PBAD*_*sfp_bpsA* | 2 |
| pTE252 | pBBR1*_oriV_aacC1_araC_PBAD_sfp_bpsA*, a gentamicin-resistant variant of pTE251 | This study |
| pJE443 | pK18mobsacB *∆ampC::PxylE-xylE:Ptac-xylAB:talB:tktA* cassette, source of *tac* promoter | 1 |
| pTE252_IND | pTE252_pBBR1*_oriV_aacC1_Ptac_sfp_bpsA* | This study |

**Supplementary Table 3. Oligonucleotides used in this study**

| **Name** | **Sequence (5’-3’)** |
| --- | --- |
| XylA_WT_F | GCCGCGCGGCAGCCATATGCAGGCCTACTTCGAC |
| XylA_WT_R | GTTAGCAGCCGGATCCTCACTTATCGAACAGGTAGTG |
| XylB_WT_F | GCCGCGCGGCAGCCATATGTACATTGGTATTGATTTGGG |
| XylB_WT_R | GTTAGCAGCCGGATCCTTAGGCCATCAGCGGC |
| XylB_G395R:1_F | CCTGATCGGCCGCGGCGCCCGGT |
| XylB_G395R:1_R | GTCACCGACTGTGGCTTGATG |
| XylB_G395W:2_F | CCTGATCGGCTGGGGCGCCCGGT |
| XylB_G395W:2_R | GTCACCGACTGTGGCTTGATGCCGC |
| XylB_recomb_ch_F | TTGGGTCCTGGTTATGCTGC |
| XylB_recomb_ch_R | CTTCCGGCAAATCGACAAGG |
| XylE_recomb_ch_F | TGCTTTTTTTCTGGCGCGTAATACAACACC |
| XylE_recomb_ch_R | AAACAAGCTTTGGGTTGTAGTGACGGAAGG |
| AraC2_recomb_ch_F | TGCTTTTTTTGGTCTGGATTCTGTGTTCGG |
| AraC2_recomb_ch_R | AAACAAGCTTTGAGAAAGTGGTGGAGGTCG |
| AraD2_recomb_ch_F | TGCTTTTTTTTGACGGTCAATGGGAAGAGC |
| AraD2_recomb_ch_R | TCTAGCTCTAAAACAAGCTTTCCAGAATCGGACGAATGCC |
| PP3603_recomb_ch_F | TGCTTTTTTTTGTGCTACCAGAACTACCCG |
| PP3603_recomb_ch_R | CTAGCTCTAAAACAAGCTTTGCAGGTGATCATAGCCAGG |
| ptac_sfp_bpsA_ins_F | TCTTATGACAACTTGACGGCTACATCATTCACGCCTGGAAGACGCAGACATTCC |
| ptac_sfp_bpsA_vec_R | GTGAATGATGTAGCCGTCAAGTTGTCATAA |
| ptac_sfp_bpsA_ins_R | TTCATATGTATATCTCCTTCTTAAAAGATCTTTTGAATTCAATTGTTATCCGCTCACAATTCCACACAT |
| ptac_sfp_bpsA_vec_F | GAATTCAAAAGATCTTTTAAGAAGGAGATATACATATGAAGATTTACG |

**Supplementary Table 4. Mutations in the starting strain JE3692**

| **Gene** | **Protein** | **Modification** | **Mutation** |
| --- | --- | --- | --- |
| PP_4387/  *flgE* (PP_4388) | Hypothetical protein/ flagellar hook protein | -GGC | Intergenic (-24/+44) |
| *rluA* (PP_1731)/ *minE* (PP_1732) | Ribosomal large subunit pseudouridine synthase/ cell division topological specificity factor | -C | Intergenic (-107/+43) |
| *xylE/ xylA* | D-xylose:proton symporter/ D-xylose isomerase | -T | Intergenic (-103/-105) |

Reference genome: *P. putida* KT2440^3^

**Supplementary Table 5. Whole-genome sequencing sample list**

| **Strains** | **Bioproject** | **Biosample** |
| --- | --- | --- |
| JE3692 | PRJNA1395279 | SAMN54354527 |
| GLC1 | PRJNA1395279 | SAMN54354528 |
| GLC2 | PRJNA1395279 | SAMN54354529 |
| GLC3 | PRJNA1395279 | SAMN54354530 |
| GLC4 | PRJNA1395279 | SAMN54354531 |
| XYL1 | PRJNA1395279 | SAMN54354532 |
| XYL2 | PRJNA1395279 | SAMN54354533 |
| XYL3 | PRJNA1395279 | SAMN54354534 |
| XYL4 | PRJNA1395279 | SAMN54354535 |
| ARA1 | PRJNA1395279 | SAMN54354536 |
| ARA2 | PRJNA1395279 | SAMN54354537 |
| ARA3 | PRJNA1395279 | SAMN54354538 |
| ARA4 | PRJNA1395279 | SAMN54354539 |
| HSM1 | PRJNA1395279 | SAMN54354540 |
| HSM2 | PRJNA1395279 | SAMN54354541 |
| HSM3 | PRJNA1395279 | SAMN54354542 |
| HSM4 | PRJNA1395279 | SAMN54354543 |
| LSM1 | PRJNA1395279 | SAMN54354544 |
| LSM2 | PRJNA1395279 | SAMN54354545 |
| LSM3 | PRJNA1395279 | SAMN54354546 |
| LSM4 | PRJNA1395279 | SAMN54354547 |
| GXA1 | PRJNA1395279 | SAMN54354548 |
| GXA2 | PRJNA1395279 | SAMN54354549 |
| GXA3 | PRJNA1395279 | SAMN54354550 |
| GXA4 | PRJNA1395279 | SAMN54354551 |
